# Tooth size apportionment, Bayesian inference, and the phylogeny of *Homo naledi*

**DOI:** 10.1101/2020.12.16.423087

**Authors:** Joel D. Irish, Mark Grabowski

## Abstract

This study has three main objectives—two methodological and one summative, namely, further characterization of *Homo naledi* (∼335–236 ka) to more firmly establish its evolutionary history. Using mathematically-corrected mesiodistal and buccolingual crown dimensions, the species was compared with samples of *Pan troglodytes, Australopithecus africanus*, *A. afarensis*, *Paranthropus robustus*, *P. boisei*, *H. habilis*, *H. ergaster*, *H. erectus*, *H. heidelbergensis*, *H. neanderthalensis*, and *H. sapiens*; the correction yields equivalently scaled samples unaffected by significant interspecific size differences. After initial cluster analysis, the data were used in tooth size apportionment analysis to determine how size is distributed relatively in each species’ dentition, while visualizing this variation in a sample scatterplot. The first main objective then, after quantitative coding, is evaluating the utility of these characters to estimate phylogenetic relationships, here using Bayesian inference with an Mkv model. The second objective, for the first time in paleoanthropological study, is estimating relationships using continuous characters, i.e., the scaled data, through Bayesian inference under a Brownian-motion model. This strategy facilitates maximum reception of potential phylogenetic signal. The final objective based on all analyses, though principally continuous Bayesian inference, is to elucidate the phylogeny of *H. naledi.* Relationships are largely congruent across methods and, with markedly higher node support, most of those inferred in prior systematic studies using qualitatively discretized traits. The present results place *H. naledi* as a sister taxon to *H. habilis* (node support ∼70-99%), with a plesiomorphic pattern of relative tooth size. It is nested within a clade comprising australopiths and early *Homo* dating 3.3 Ma to ∼800 ka, distinct from younger *H. erectus* through *H. sapiens.* This suggests that *H. naledi* originated well before the geological date range associated with the Dinaledi Chamber, from which the remains in this study were recovered, to represent a long-lived side branch in the genus.

## 1. Introduction

In this study tooth size apportionment (TSA) analysis was used to characterize further the recently defined hominin, *Homo naledi* (Berger et al., 2015), ∼335–236 ka (Dirks et al., 2017), relative to other Plio-Pleistocene and recent comparative species. It builds on work demonstrating the efficacy of this odontometric technique for gauging phenetic affinities among various African hominin samples, including *Australopithecus sediba*, along with recent humans and *Pan troglodytes* (Irish et al., 2016). In TSA, the unit of study is the entire permanent dentition, rather than individual mesiodistal (MD) and buccolingual (BL) crown dimensions. That is, these lengths and widths are corrected mathematically (below) to yield equivalently scaled samples, which are then submitted to principal components analysis to produce statistically uncorrelated factor scores; these scores are compared to determine how crown size is differentially distributed, or apportioned, along the tooth rows. Because TSA was found to be effective in comparing both modern human individuals and groups (Harris and Bailit, 1988; Harris and Rathbun, 1991; Hemphill et al., 1992; Lukacs and Hemphill, 1993; Harris, 1997, 1998; Irish and Hemphill, 2001; Irish and Kenyhercz, 2013), which on an intraspecific level exhibit minimal variation, the approach was projected to be particularly effective in comparing more discernible interspecific dental differences in our hominin ancestors. This prediction was proven correct, as the grouping of species (in Irish et al., 2016) that were also included in other, albeit, cladistic studies (Strait et al., 1997; Strait and Grine, 2004; Smith and Grine, 2008) is congruent, as are results pertaining to specific relationships of *A. sediba* (Berger et al., 2010; Irish et al., 2013, 2014; Dembo et al., 2015, 2016).

As such, the initial intent here was to simply undertake equivalent TSA analyses to assess the interspecific relationships of *H. naledi*, as well as its taxonomic classification. Earlier studies using characters throughout the skeleton, including dental nonmetric traits, supported its inclusion in the genus *Homo*, but as a distinct member (Berger et al., 2015; Thackeray, 2015; Dembo et al., 2016; Irish et al., 2018; also Holloway et al., 2018; Davies et al., 2020). A phenetic approach was deemed most appropriate because odontometric data, original or scaled, do not lend themselves well to cladistics. Of course, the same caveat applies to all continuous morphological data in traditional, non-probabilistic phylogenetic analyses (Pimentel and Riggins, 1987; Pogue and Mickevich, 1990; Thiele, 1993; Wiens, 1995; Poe and Wiens, 2000; Parins-Fukuchi, 2018a), when not excluded entirely (Poe and Wiens, 2000; Garcia-Cruz and Sosa; 2006). Defined, these are data that “… can take on any real number value, whether a measurement of a morphometric character in a specific individual, or the mean value of an intraspecifically variable, quantitative trait for a given species” (Wiens, 2001:690).

Standard operating procedure to include such data in cladistic analyses necessitates subjective, qualitative coding (per Stevens, 1991; Thiele, 1993; Wiens, 2001; Felsenstein, 2004; Parins-Fukuchi, 2018a, 2018b) into two or a few discrete states, or by using one of several more objective, quantitative techniques. Some of the latter, to boost phylogenetic signal over qualitative discretizing, return upwards of 30+ character states (reviewed in Wiens, 2001; Felsenstein, 2004; Garcia-Cruz and Sosa; 2006; Jones and Butler, 2018), including oft-used gap-weighting (Thiele, 1993; Schols et al, 2004; Garcia-Cruz and Sosa; 2006; Goloboff et al., 2006). Yet, quantitative coding comes with other concerns, and still fails to detect all key information (Farris, 1990; Wiens, 2001; Felsenstein, 2004; Goloboff et al., 2006; Parins-Fukuchi, 2018b).

So, while coded data remain standard for phylogenetic analyses of fossil hominins (e.g., Strait et al., 1997; Strait and Grine, 2001; 2004; Dembo et al., 2015, 2016; Mongle et al., 2019), a relatively rare yet promising alternative (Parins-Fukuchi, 2018a, 2018b) is to directly analyze continuous characters (Smith and Hendricks, 2013). Specifically, advances in probabilistic phylogenetics, notably Bayesian inference, allow the use of models capable of approximating the evolution of continuous morphological characters (Felsenstein, 2004; Höhna et al., 2016; Parins-Fukuchi, 2018a, 2018b). Among other strengths (see below), the latter are more objectively collected through standardized measurements, do not require the ordering of states, and retain phylogenetic information at much higher evolutionary rates than do coded characters—precisely because they do not necessitate potentially subjective ‘compression’ into a finite number of states (Parins-Fukuchi, 2018a, 2018b).

Thus, analyses of *H. naledi*, plus nine comparative samples of African and Eurasian Plio-Pleistocene hominin species, two regional samples of recent African *H. sapiens*, and *Pan troglodytes* will proceed as follows. First, as before (Irish et al., 2016), a standard phenetic strategy, using cluster analysis, principal components analysis, and a scatterplot is followed to obtain interspecific affinities from the continuous scaled MD and BL data. Second, for the first time in the study of hominin taxa, phylogenies are inferred based on continuous data using a Brownian-motion model and Bayesian inference (Höhna et al., 2016; Parins-Fukuchi, 2018a). For methodological comparison, relationships will initially be estimated via a more common Bayesian strategy, using quantitatively-coded versions of the scaled dimensions under an Mkv (Lewis, 2001) or “standard discrete (morphology)” model (Huelsenbeck and Ronquist, 2001; Ronquist and Huelsenbeck, 2003; Ronquist et al., 2020:133). This extended analytical approach also serves to explore further the effectiveness of these odontometric characters for inferring hominin phylogenies. Finally, the results will be compared across all methods for similarity, or the lack thereof, and in sum with those from the abovementioned studies to provide additional characterization of *Homo naledi*.

## 2. Materials and methods

### 2.1 The samples and their data sources

The *H. naledi* sample consists of 122 dental specimens from the Dinaledi Chamber of the Rising Star cave system, South Africa, with mean MD and BL measurements used in the present analyses from Berger et al. (2015). These fossils are directly linked to the ∼335–236 ka geological age (Dirks et al., 2017). The nine comparative Plio-Pleistocene samples were then chosen based on two criteria. First, they provide a cross section of the three principal later hominin genera, although with an emphasis on *Homo*. Second, while an analysis of individual hominins can be approximated (Irish et al., 2016), all samples, like *H. naledi*, have multiple MD and BL measurements for the permanent teeth; this yields the most accurate species means to reduce issues related to very small sample sizes. Therefore, *Kenyanthropus platyops, A. sediba, P. aethiopicus, H. rudolfensis, and H. floresiensis,* among other species, were not included. Mean data for six samples, all African, are the same as those published in the prior TSA study (Irish et al., 2016), with source information included therein. They are *A. africanus* (*n=*307 total teeth), *A. afarensis* (*n=*271), *Paranthropus robustus* (*n=*377), *P. boisei* (*n=*172), *H. habilis* (*n=*93), and *H. ergaster* (*n=*260). Because few anterior teeth for the latter species have been found in Africa, MD and BL means again include published dimensions in 38 crowns from Dmanisi attributed to *H. ergaster* (following Martinón-Torres et al., 2008; Rightmire and Lordkipanidze, 2010; Lordkipanidze et al., 2013; among others). The three non-African samples include *H. erectus* (*n=*587) (data compiled for the present study from Weidenreich, 1937, 1945; Wu and Chia, 1954; Jacob, 1973; Bermudez de Castro, 1986; Wood, 1991; Wu and Poirier, 1995; Kaifu et al., 2005; Zaim et al., 2011; Xing et al., 2018), *H. heidelbergensis* (*n=*789), and *H. neanderthalensis* (*n*=821 teeth). The MD and BL mean data for the latter two are from Berger et al. (2015), where source information is listed.

Representing *H. sapiens* are two large regional samples comprising post-Pleistocene North (*n*=20,674 teeth, 1412 individuals) and sub-Saharan Africans (*n*=15,948, 822 inds.), directly recorded by the first author (sample details in Irish, 1993, 2000, 2005, 2006, 2008, 2010). Finally, to emphasize among-species taxonomic variation (Mahler, 1973) and help illustrate methodological effectiveness in the phenetic analyses, while also serving as the root for the phylogenetic inference (below), the same *Pan troglodytes* data (*n*=924 teeth, 70 individuals) used previously are included (Irish et al., 2016).

### 2.2 Odontometric measurements

For the two African *H. sapiens* samples, maximum dimensions were recorded in each tooth following established protocol (Moorrees and Reed, 1964). The MD measurement was taken parallel to the occlusal and labial/buccal surfaces of the crown; the BL was measured as the maximum distance perpendicular to the MD (Hemphill, 2016a). Needlepoint callipers accurate to 0.05 mm were used. If antimeric pairs in individuals were present, then mean MD and BL data were calculated for the analysis as possible; if only the right or left tooth remained, its measurements were used, for up to 16 measurements in each isomere and 32 per dentition. For all other samples, measurement techniques were reviewed for conformity to facilitate data compatibility, though inter-observer error obviously could not be tested.

Promoting credible representative affinities, the narrow-sense heritability of MD and BL diameters is high, in some cases *h*^2^>0.8 (Alvesalo and Tigerstedt, 1974; Townsend and Brown, 1980; Kieser, 1990; Dempsey et al., 1995; Dempsey and Townsend, 2001; Hlusko et al., 2002; Townsend et al., 2003; Baydas et al., 2005; Rizk et al., 2008). A recent study of just MD diameters reports a lower mean estimate of *h*^2^=0.51, though reproductive isolation and socioeconomic stress in the population sampled and some small samples likely affected the value (per Stojanowski et al., 2017). Heritability of the mathematically corrected dental data in the present study has not been assessed directly, but should be analogous to the original MD and BL dimensions (above) given the high correlation between these two sets of data (*r*=0.93, *p*=0.00). At any rate, heritability of 0.5 to >0.8 in the Plio-Pleistocene species cannot be asserted. However, the prospect is encouraging for assessing phylogenetic relationships derived from two basic, readily available measurements.

### 2.3 Tooth size apportionment analysis

TSA analysis entails submitting a correlation matrix of data to principal components analysis (PCA), with the resulting uncorrelated components used to identify patterning of inter-tooth relationships. However, because this study is inter-rather than intraspecific, the methodology of previous TSA research (Harris and Bailit, 1988; Harris and Rathbun, 1991; Hemphill et al., 1992; Lukacs and Hemphill, 1993; Harris, 1997, 1998; Irish and Hemphill, 2001; Hemphill, 2016b) was tailored to address the substantial tooth size differences, e.g., *Paranthropus* vs. *Pan* (Irish et al., 2016). Like all skeletal measurements, odontometric data can be split into two components: 1) (absolute) size and 2) shape (relative size) (Penrose, 1954; Rahman, 1962; Corruccini, 1973; Harris and Harris, 2007; Townsend et al., 2009; Irish et al., 2016). So a mathematical correction presented in Jungers et al. (1995:145) that they termed “DM_RAW,” from Darroch and Mosimann, (1985), was used to minimize size effects that dominate the first principal component, contra residual scores commonly substituted for modern humans (e.g., Harris, 1997). The geometric mean (GM) is computed as the *n*th root of the product for all *n* dimensions (x) per case. Each dimension is then divided by this mean (x/GM) for that case to provide an average of 1.0 across each sample row. This scaling “cancels out size differences by giving each [sample] the same average character state or magnitude over all the measurements taken on it” (Corruccini, 1973:747).

Prior to PCA, basic descriptive results were calculated and sample data submitted to hierarchical agglomerative cluster analysis, based on Euclidean distances; in this way, initial impressions of the relationships could be obtained. The average linkage (between groups), a.k.a. UPGMA (unweighted pair group method with arithmetic mean) algorithm was applied to produce the dendrograms (e.g., Sokal and Sneath, 1963; Everitt, 1980; Aldenderfer and Blashfield, 1984; Romesburg, 1984; Felsenstein, 2004). UPGMA starts by considering each sample as an individual cluster. The two most similar clusters, consisting of 1-*n* samples, are then combined based on the smallest average inter-sample distance between them. The process continues until a single cluster results. Then, for the TSA analysis, the correlation matrix of 32 DM_RAW-scaled mean MD and BL diameters was submitted to PCA, with the resulting group component scores plotted to visualize further the phenetic variation. The program PAST 4.03 (Hammer et al., 2001) was used for cluster analysis, FigTree 1.1.4 for the dendrograms, and SPSS Ver. 26.0 for the PCA and sample plotting in three dimensions.

TSA analyses would ideally be conducted with samples divided by sex, although this strategy was not followed in the aforementioned comparisons of modern humans. That is, while sexual dimorphism may be a factor in absolute crown size differences between males and females from the same population (though see Harris, 2003), relative apportionment of tooth size within dentitions is not affected (Harris and Rathbun, 1991; Hemphill et al., 1992; Hemphill, 2016b). As with heritability, the same cannot be claimed for the Plio-Pleistocene species, or for that matter *Pan troglodytes*, with major body size differences between males and females. In any event, it is out of necessity, including an inability to assign sex to most hominin specimens, considerable missing data, and a need to maximize sample sizes, that all specimens and individuals were pooled according to species for analysis.

### 2.4 Bayesian phylogenetic inference with quantitatively coded odontometric data

Probabilistic or statistical methods to infer phylogenies, which include Bayesian inference, as well as maximum likelihood, are seeing increased usage over non-probabilistic methods such as maximum parsimony. The reasons include methodological consistency, the ability to estimate branch lengths and evolutionary rates and, basically, better performance of the former two methods in both genetic and morphological cases (Felsenstein, 2004; Lee et al., 2014; Wright and Hillis, 2014; EC.Europa.EU, 2016; Nascimento et al., 2017; Parins-Fukuchi, 2018a, 2018b; Guillerme and Brazeau, 2018). Indeed, the “… inconsistency of parsimony has been the strongest challenge to its use,” although the method is considered well suited for very large datasets, particularly of recently derived species (Felsenstein, 2004:121; EC.Europa.EU, 2016).

Of the two probabilistic methods, most recent studies applied Bayesian inference, due to the availability of programs, simpler principles relative to maximum likelihood and, arguably, more appropriate models (Mk and Mkv) for use with morphological characters (Huelsenbeck and Ronquist, 2001; Lewis, 2001; Ronquist and Huelsenbeck, 2003; Pagel and Meade, 2004; Drummond and Bouckaert, 2014; Goloboff and Catalano, 2016; Nascimento et al., 2017; Parins-Fukuchi, 2018a, 2018b; Ronquist et al., 2020). These reasons encourage application, but in this study it is also because, with an appropriate model, the phylogenetic method can handle continuous character data (Höhna et al., 2016; Parins-Fukuchi, 2018a; detailed below). The theories behind, overviews of, and techniques concerning Bayesian inference for parameter estimation are covered comprehensively in the above references, and have been discussed in prior hominin studies (Dembo et al., 2016; Mongle et al., 2019). Here, additional information pertinent to paleoanthropological research is provided during the course of describing the analytical progression.

For purposes of methodological comparison with the results from continuous data, phylogenies were initially inferred using quantitatively coded versions of DM_RAW-scaled data with an Mkv or standard discrete model (Huelsenbeck and Ronquist, 2001; Ronquist and Huelsenbeck, 2003; Ronquist et al., 2020). The 32 scaled dimensions were gap-weighted using Thiele’s (1993) method in MorphoCode 1.1, which can calculate a range of states from three to 26 (Schols et al., 2004). It generates a data matrix, in which the order and dispersal of means are determined for each morphological character, and converted to “ordered, multistate characters where the distance between means is represented by the distance between ordered character states” (Thiele, 1993; Wiens, 2001; Schols et al., 2004:2). This matrix, in Nexus format, was submitted to MrBayes 3.2.7 (Huelsenbeck and Ronquist, 2001; Ronquist and Huelsenbeck, 2003; Ronquist et al., 2020) using the maximum states allowed by the latter program (see below).

Given the vast range of parameters in MrBayes pertaining to topology, branch lengths, evolutionary rates, and others, the aim was to begin simply, using a rooted strict-clock model with default, “so-called flat, uninformative, or vague [prior] distributions” (Felsenstein, 2004; EC.Europa.EU; 2016; Ronquist et al., 2020:91). The latter are suggested to base the posterior distributions principally on the data—to establish their effectiveness (Ronquist et al., 2020; though see Felsenstein, 2004; Nascimento et al., 2017). From this starting point, increasingly informative alternative parameters were added in a series of analyses. Of these, two rooted relaxed-clock models are presented as representative of this progression: one basic and the other incorporating numerous constraints, calibrations, and additional parameters. All entail Bayesian molecular clock methods to estimate divergence among the taxa (Hedges and Kumar, 2009; Nascimento et al., 2017).

As standard in Bayesian inference, each model was analyzed via Markov chain Monte Carlo (MCMC) simulation with the Metropolis algorithm (Felsenstein, 2004; EC.Europa.EU; 2016; Nascimento et al., 2017; Ronquist et al., 2020). Because the dataset is not very large, the MrBayes default run values were used, with an increase in generations as necessary. Two simultaneous but independent analyses beginning with different random trees were run for 1,000,000 generations, with a sampling frequency of 500 to yield 2000 samples, and diagnostics calculated every 5000 generations. Each run consisted of one cold and three heated chains, with a 25% burn-in of samples from the cold chain to settle into its equilibrium distribution. In this way, expedient calculation of convergence diagnostics is possible to assess if a representative sample of trees resulted from the posterior probability distribution.

Established diagnostics used for the coded data include: 1) standard deviation of split frequencies ≤0.01, 2) potential scale reduction factor (PSRF) of ∼1.0 for all parameters, and 3) average effective sample sizes (ESS) of >100, but aiming for >200 (Huelsenbeck and Ronquist, 2001; Ronquist and Huelsenbeck, 2003; Felsenstein, 2004; EC.Europa.EU, 2016; Nascimento et al., 2017; Guillerme and Brazeau, 2018; Ronquist et al., 2020). If any cut-offs were not met, the generation number was increased until minimums were achieved or exceeded, to result in similar trees from the independent runs. Finally, a cladogram with posterior probabilities, a.k.a. clade credibility values for all internal nodes, and a phylogram of the mean branch lengths were produced. FigTree 1.1.4 was again used to render trees. Associated diagnostics include posterior probabilities to indicate final tree number where, for example, a value of ∼1.0 provides support for a single tree (Ronquist et al., 2020).

The three clock analyses described here share several initial priors particular to the quantitatively coded data type [full parameter list in Supplementary Information (SI) Table S1]. For the state frequencies a symmetric dirichlet distribution fixed to infinity was used to correspond to the assumption of no transition rate asymmetry across sites; the rate from 0 to 1 is equal to that for 1 to 0 for each character (Ronquist et al., 2020). The coding bias was variable and type ordered, as necessary for the gap-weighted continuous data, where it is assumed that the evolution between states moves through intermediate states (0 <-> 1 <-> (Felsenstein, 2004). MrBayes can handle 10 character states (0-9) if unordered, but just a maximum of six when ordered (Ronquist et al., 2020). Thus, only state values of 0-5 were calculated in Morphocode 1.1 for the Nexus input matrix.

First, for the strict-clock analysis (SI Table S1), a clock parameter was specified for branch lengths type, with a uniform prior [i.e., tree age with gamma distribution, node ages not constrained], and fixed clock rate. Thus, constant rates of evolution are assumed among the taxa, where all branch tips are presumed to be the same age (Felsenstein, 2004; Pybus, 2006; EC.Europa.EU, 2016; Ronquist et al., 2020). This somewhat restrictive approach is favored for analyses of the same species with similar molecular evolution rates (Felsenstein, 2004), which may not be the case for the present hominin taxa.

Second, a basic relaxed-clock analysis (SI Table S1) was conducted. Following on from above, a model of this type is suggested for biologically distinct taxa (i.e., different species), because it can incorporate a prior distribution of evolutionary rates that vary among taxa and branches of a phylogeny (Felsenstein, 2004; Pybus, 2006). A relaxed-clock model has more parameters than a strict-clock, but is again rooted. That said, the information on root position is weaker, so following advised protocol a tree topology constraint was introduced to exclude *Pan* and force all hominin taxa into a monophyletic ingroup. The key change was to “’relax’ the strict clock assumption” with an independent gamma rates (IGR) model of continuous uncorrelated variation across lineages (Ronquist et al., 2020:60). A related prior was also added for a standard exponential rate of variance in effective branch lengths over time. Lastly, a gamma distribution was substituted for the rates prior (also in Dembo et al., 2016). It is the simplest, most common non-default prior to accommodate rate variation across sites (Kuhner and Felsenstein, 1994; EC. Europa.EU, 2016; Ronquist et al., 2020). The other parameters were the same as above.

Third, a second relaxed-clock analysis (see SI Table S1) incorporated two important additions: 1) calibration node dating with a uniform prior of 5-6 Ma for the root, *Pan*, to designate the likely chimp/hominin split, in combination with 2) tip dating, where the node depths are constrained by calibrating the hominins using a uniform prior with minimum and maximum ages; thus, estimated extinction and origination dates (after Du et al., 2020) were included: *A. africanus* (2.3, 3.3 Ma), *A. afarensis* (2.95, 3.24), *P. robustus* (1.6, 2.36), *P. boisei* (1.41, 1.93), *H. habilis* (1.15, 2.038), *H. ergaster* (0.79, 1.945), *H. erectus* (0.143, 1.81), *H. heidelbergensis* (0.1, 0.99) (Du et al., 2020), *H. neanderthalensis* (0.028, 0.2) (Wood and Lonergan, 2008), and *H. sapiens* (0.0, 0.315) (Richter et al., 2017). For *H. naledi* the date from Dirks et al. (2017) (i.e., 0.236, 0.335) was used, with the caveat that it applies to the Dinaledi chamber and not necessarily the full temporal range of the species.

Three notable changes were also made to this dated relaxed-clock model. First, the clock rate default prior, which measures node age as the number of expected substitutions per site, was swapped for a normal distribution to calibrate the tree in millions of years. A mean of 0.2 and standard deviation of 0.02 designated this distribution to be truncated at 0 to yield positive values (Ronquist and Huelsenbeck, 2003; Ronquist et al., 2020). Second, the gamma rates parameter was replaced with an alternative that produces equivalent results— the lognormal distribution (Olsen, 1988; Kuhner and Felsenstein, 1994; Thorne et al., 1998; Kishino et al., 2001). The change was made, simply, to promote methodological comparison, as lognormal is a suggested prior (in Heath, 2019) for RevBayes 1.1.0 (Höhna et al., 2016), the program employed to infer phylogenies from continuous DM_RAW-scaled data. Third, a fossilized birth-death (FBD) prior with random sampling was substituted for the uniform branch lengths default (Ronquist, et al., 2020).

Oftentimes a standard birth-death prior (e.g., Dembo et al., 2016) is used with dating and root constraints (Nascimento et al., 2017; Ronquist et al., 2020). However, the FBD is most appropriate for clock trees with calibrated external nodes (i.e., fossils) and if samples of both extinct and extant species are included (Stadler, 2010; Heath et al., 2014; Heath, 2015; Stadler et al., 2018; Zhang et al., 2015; Ronquist, et al., 2020). This prior describes the probability of tree and fossil data conditional on several birth-death parameters (SI Table S1), including speciation with branching (birth), extinction (death), and fossil preservation and recovery (sampling). Most importantly, the FBD allows for the inclusion of stratigraphic ranges of fossil taxa, i.e., the above dates, to accommodate for the conclusion that ignoring uncertainty in age estimates can yield bias in divergence times (Barido-Sottani et al., 2019).

Finally, all clock models were compared by calculating Bayes factors (B_12_) from the marginal likelihoods that result when substituting the MCMC with the stepping-stone (ss) method (Xie et al., 2011). To do so in MrBayes, it is suggested that the MCMC generation number be increased by a factor of 10 (Ronquist et al., 2020). This ss method, though with modification, was also used for the analysis of continuous data (see below). For both, the difference between logarithms of the marginal likelihoods was doubled [2*log*_e_*(B_12_)] where, based on proven criteria, a product between 0 and 2 indicates “not worth more than a bare mention,” 2-6 is “positive,” 6-10 “strong,” and >10 “very strong” evidence either against or for model 1 (M_1_) vs. model 2 (M_2_) (Kass and Raftery, 1993, 1995:777; Ronquist et al., 2020).

### 2.5 Bayesian phylogenetic inference with continuous odontometric data

Continuous data can be treated differently from coded characters for phylogenetic inference by choosing a specific model of evolution for how traits evolve on the tree. One of the most common is Brownian-motion (Felsenstein, 1985, 2004). It is a fairly simplistic model (Parins-Fukuchi, 2018b) often used to approximate evolution under drift or adaption to a randomly fluctuating optimum (Felsenstein, 1985, 2004; Hansen and Martins, 1996). Under Brownian-motion, evolution is described as a random walk, where expected changes in traits along branches of the tree have an expected value of 0 and a variance proportional to branch length (Felsenstein, 1985; Revell et al., 2008; Parins-Fukuchi, 2018a). Brownian-motion has been used a number of times to successfully model dental evolution on a phylogeny (e.g., Gomez-Robles et al., 2013), including in simulations where the underlying model used to generate the “true” phylogeny was not Brownian-motion (per Parins-Fukuchi, 2018a). Following previous work, this assumption was made here.

For this study, phylogenetic inferences were based on actual DM-scaled dimensions with RevBayes (Höhna et al., 2016). Like its predecessor MrBayes, RevBayes uses MCMC to compute joint posterior probability distribution of model parameters. The basic structure presented by Parins-Fukuchi (2018a) was followed for the continuous odontometric data, but the present approach was set up to mirror the analyses described for the quantitatively-coded data: 1) a strict-clock model, 2) a basic relaxed-clock model, and 3) a relaxed-clock model incorporating the abovementioned fossil date ranges to calibrate the terminal nodes. For each model, two MCMC simulations were run for 2,500,000 to 5,000,000 generations, with a burn-in of 50,000 generations and a tuning interval of 100 generations. Generation time was set so that effective sample size of all parameters exceeded 200 (following Parins-Fukuchi, 2018a). Chain mixing was confirmed visually with traceplots in Tracer 1.7.0 (after Parins-Fukuchi, 2018a). Maximum *a posteriori* trees showing a topology with the highest posterior probability and mean of the posterior branch distribution were calculated based on the two chains, and are included in the results section with the node support shown. Maximum *a posteriori* trees are interpreted as those most probable to have generated the observed data (Höhna et al., 2017a).

Though the three coded analyses setups were emulated, some modifications were necessary because of differences associated with continuous data, such as the assumption of Brownian motion for the model. The continuous strict-clock setup used a prior uniform distribution on the clock rate, an exponential distribution on root age, a standard prior on the Brownian-motion rate parameter (sigma, or log sigma for efficiency in inference), and a uniform prior where equal probability is placed on different topologies and node ages (SI Table S2). This prior allows the continuous data themselves to provide the branch length information (Höhna et al., 2017b). The basic relaxed-clock setup (SI Table S2) builds on the strict-clock model but, following above, uses an uncorrelated gamma distributed prior on the branch rates. Here, the rate associated with each branch is a stochastic node, which is the combination of an independent draw from a gamma distribution and an exponentially distributed global clock rate (Kuhner and Felsenstein, 1994; EC. Europa.EU, 2016; Ronquist et al., 2020).

The third and final continuous data analysis used the fossilized birth-death (FBD) model as already described (Heath et al., 2014; Stadler et al., 2018; Höhna et al., 2016), and included the same stratigraphic date ranges previously listed for all taxa. Again, the rate associated with each branch is the combination of an independent draw, although now from a lognormal distribution (above), and an exponentially distributed global clock rate (Olsen, 1988; Kuhner and Felsenstein, 1994; Thorne et al., 1998; Kishino et al., 2001; Heath, 2019). The other parameters were largely similar to those of the continuous basic relaxed-clock model, but key changes included a uniform prior set on the tree origin time, an exponential prior on fossil sampling rate, an exponential prior on speciation rate, and a beta prior on extinction rate (SI Table S2). In addition, while the continuous basic relaxed-clock analysis did not need it, this FBD model required a constraint to exclude the root, *Pan*, from the hominin ingroup. In this case, exclusion was achieved by forcing the root to become ‘extinct’ before origination of the hominins, using the 5-6 Ma range from the coded dated relaxed-clock analysis. And again, model selection between this and the other continuous analyses was performed via stepping-stone sampling to estimate marginal likelihoods to calculate Bayes factors (Xie et al., 2011), though using the protocol suggested for RevBayes (Höhna et al., 2016, 2017a; and below).

### 2.6 Phylogenetics and character independence

Before proceeding, some discussion about correlation among characters is in order. Phylogenetic analyses, whether probabilistic or other, assume character independence; if violated, where characters provide redundant information, the expectation is that results may be biased (e.g., Farris, 1983; Kluge, 1989; O’Leary and Geisler, 1999; Felsenstein, 2004; Strait and Grine, 2004; Gomez-Robles and Polly, 2012; Klingenberg, 2014; Kay, 2015; Parins-Fukuchi 2018a; Billet and Bardin, 2018; Guillerme and Brazeau, 2018). In TSA analysis any statistical correlation of continuous data is negated with PCA, e.g., Pearson correlations of MD and BL dimensions across the present samples revealed an *r* ≥ 0.5 in 23.5% of the 496 total pairwise comparisons. However, this does not address evolutionary correlation said to predominantly affect morphological characters (though see below), whether qualitatively vs. quantitatively coded or continuous (Wiens, 2001). It may result from such factors as genetic linkage, similar selection pressures, pleiotropy, and structural and/or organismal integration (O’Leary and Geisler, 1999; Maddison, 2000; Felsenstein, 2004, Strait and Grine, 2004; Hlusko and Mahaney, 2009; Gomez-Robles and Polly, 2012; Adams and Felice 2014; Klingenberg, 2014; Guillerme and Brazeau, 2018). A potential non-evolutionary source is character choice and coding (descriptively redundant, describing different parts of the same character, etc.) (Guillerme and Brazeau, 2018).

Coding correlation can be more readily addressed (Strait and Grine, 2004; Guillerme and Brazeau, 2018), and is not a factor with non-coded continuous characters. Evolutionary correlation—of equal concern if the data are coded or continuous (Parins-Fukuchi, 2018a, 2018b), may be studied *a posteriori* through use of phylogenetic hypotheses (Guillerme and Brazeau, 2018). Otherwise, with exception (below), effects of correlation on phylogenetic inference cannot be verified or addressed, especially when the focus of study is fossil taxa (see O’Leary and Geisler, 1999; Felsenstein, 2004; Billet and Bardin, 2018; Guillerme and Brazeau, 2018). For that matter, the same goes for possible homoplasy, another effect said to be inherent with the use of morphological data—that, nonetheless, are a necessity in the analysis of fossils (Wiens, 2001; Kay, 2015).

One likely source of evolutionary correlation that has been addressed, and is of particular relevance here, is morphological integration—as in serially homologous structures like teeth (Strait and Grine, 2004; Gomez-Robles and Polly, 2012; Klingenberg, 2014; Billet and Bardin, 2018). For example, Gomez-Robles and Polly (2012) found integration to affect the entire post-canine dentition regarding crown shape. Certain intra-oral regions and individual teeth are more strongly integrated than others, including the entire mandibular dentition (relative to maxillary), molars (vs. premolars), and the UM3 and LP3. According to the authors, such a strong departure from neutral evolution can bias phylogenetic inference and decrease “the ability to correctly classify hominin specimens and species” (Gomez-Robles and Polly, 2012: 1033). One counter measure, according to Billet and Bardin (2018), is to merge multiple dental observations into a single character.

On the other hand, the necessity and advantages of dental data in characterizing and comparing fossil taxa are well known, as noted (also Gomez-Robles and Polly, 2012), while merging dental characters risks unverified *a priori* dismissal of phylogenetic signal (O’Leary and Geisler, 1999). In fact, the most substantial size variance occurs among the tooth types (Harris, 2003), while incisors were found to be genetically independent from the post-canine dentition—at least in baboons (Hlusko and Mahaney, 2009), and studies based on separate, potentially integrated dental data have provided reliable results (e.g., O’Leary and Geisler, 1999; O’Leary et al., 2013; Kay, 2015). Along these lines, and contrary to above, evolutionary correlation, including structural integration of the dentition, is actually suggested by some workers to have relatively minimal influence on phylogenetic reconstruction (Adams and Felice, 2014; Parins-Fukuchi, 2018a). And, in reality, it is to be expected in and indicative of species that are closely related (Felsenstein, 1985; Martins and Hansen, 1997; Lajeunesse and Fox, 2015).

Whatever the case, quantitative data (coded or continuous) are no more susceptible to correlation than qualitatively coded characters that have traditionally been employed in phylogenetic analyses (per Wiens, 2001; Parins-Fukuchi, 2018a). Moreover, it is likely that molecular data are also affected, but “… phylogenetic analyses that do not accommodate for [correlation] are routinely applied” (Parins-Fukuchi, 2018a:330). Any prospective issues with the present data are also mitigated somewhat by the use of Bayesian inference, which is said to be consistently less affected than maximum parsimony, notably for comparing a small number of taxa (Guillerme and Brazeau, 2018). Brownian motion is particularly effective, relative to more complex models (Butler and King, 2004; Gomez-Robles and Polly, 2012; Lajeunesse and Fox, 2015), and when using a moderate amount of continuous data (Parins-Fukuchi, 2018a, 2018b). Beyond this, little can be done about correlation other than consider the plausibility, or lack thereof, of the resulting phylogenies *a posteriori*.

## 3. Results

### 3.1 Descriptive and tooth size apportionment analyses

The mean MD and BL dimensions of *H. naledi* and the 12 other samples, with total number of teeth from which each were calculated, are provided in Table 1. Maxillary and mandibular crown surface areas (MD x BL) were also determined and plotted (Figs 1-2). Though crude estimates of actual areas (Garn et al., 1977; Hemphill, 2016a), these products are useful indicators of absolute dental size variation among species. For both isomeres, *H. naledi* is in the bottom half of the graphs with others of its assigned genus, approximately halfway between the small-toothed *H. sapiens* and larger-toothed *H. habilis*. *Homo naledi* incisors and canines are comparable in crown area to those of *H. sapiens*, while its posterior teeth, especially P3, P4, M2, and M3 trend more toward other the *Homo* species, with the exception of *H. habilis* (compare individual measurements in Table 1).

**Table 1.**
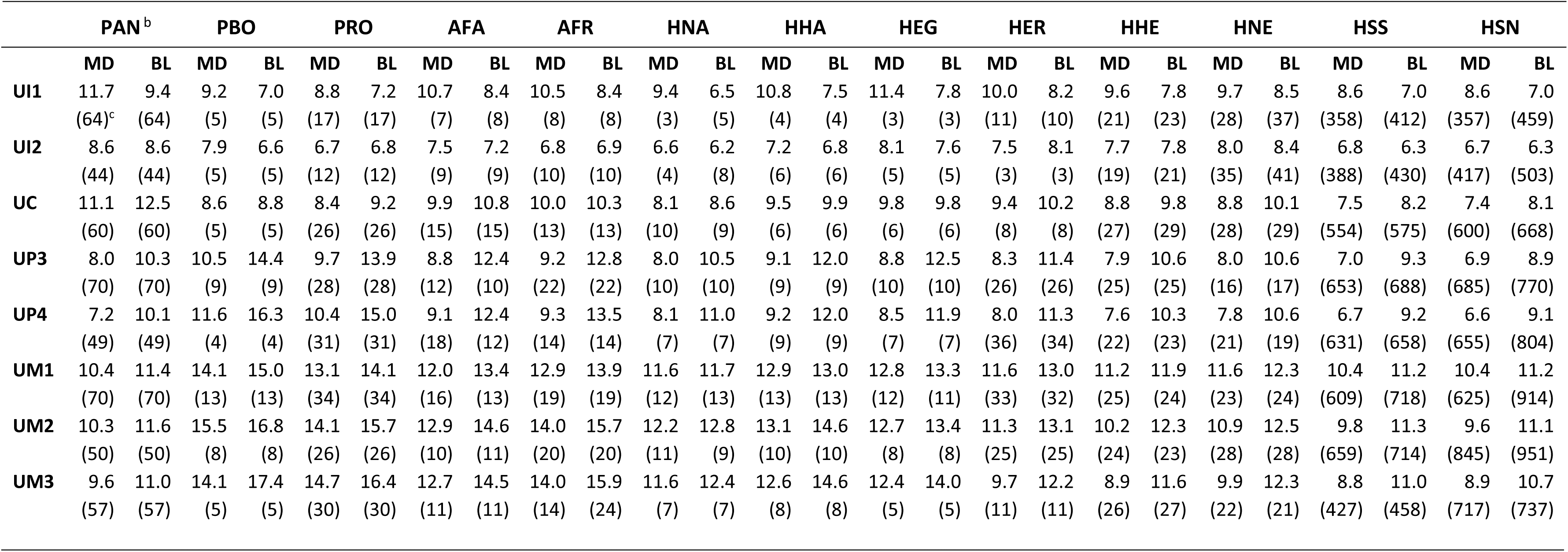
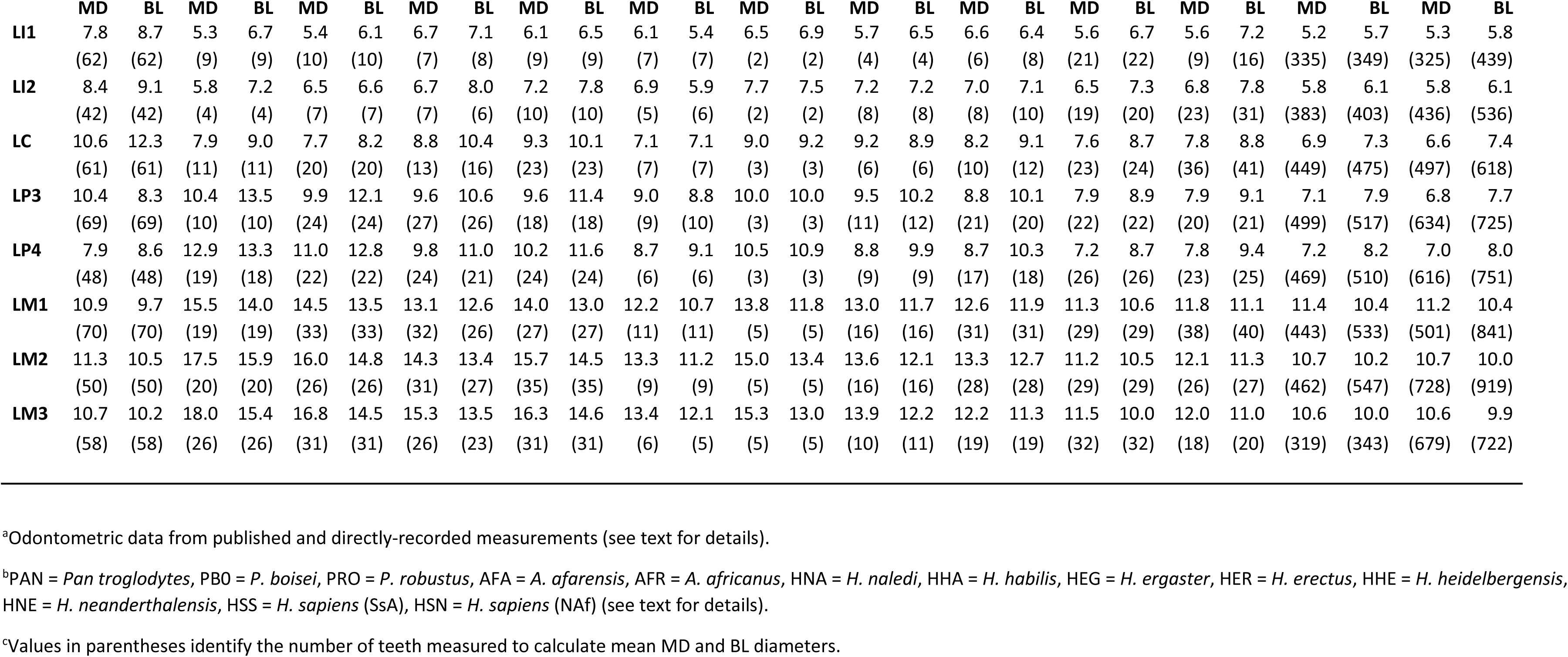
Mean maxillary and mandibular mesiodistal (MD) and buccolingual (BL) diameters^a^ for the 13 samples.

**Figure 1.**
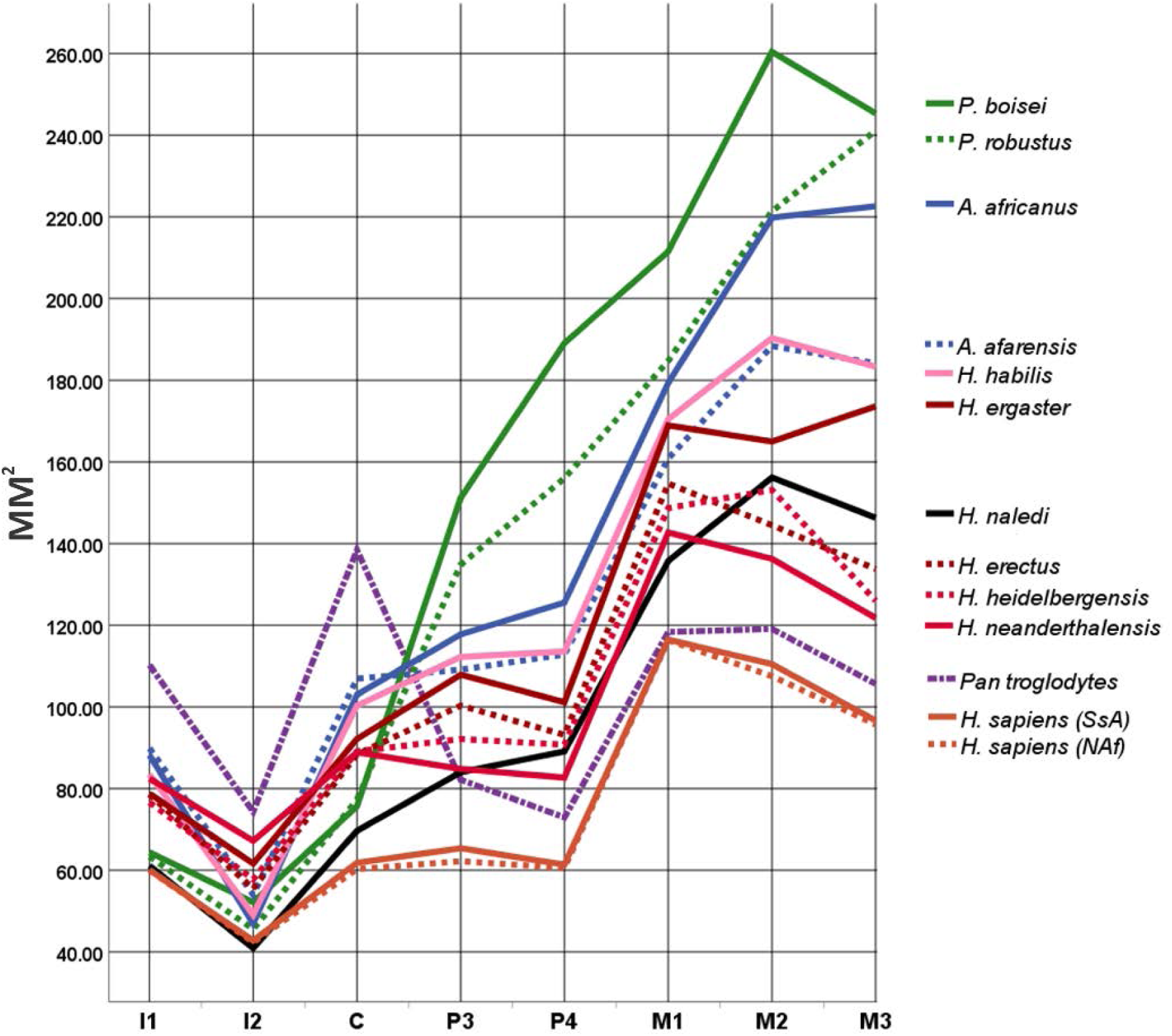
Line plot showing tooth-by-tooth trends in absolute occlusal surface areas of the maxillary dentition in mm^2^ by sample. Line colors and format apply loosely to genus (e.g., see Fig. 5, SI Fig. S1), but are primarily used to differentiate samples. See text for sample compositions.

**Figure 2.**
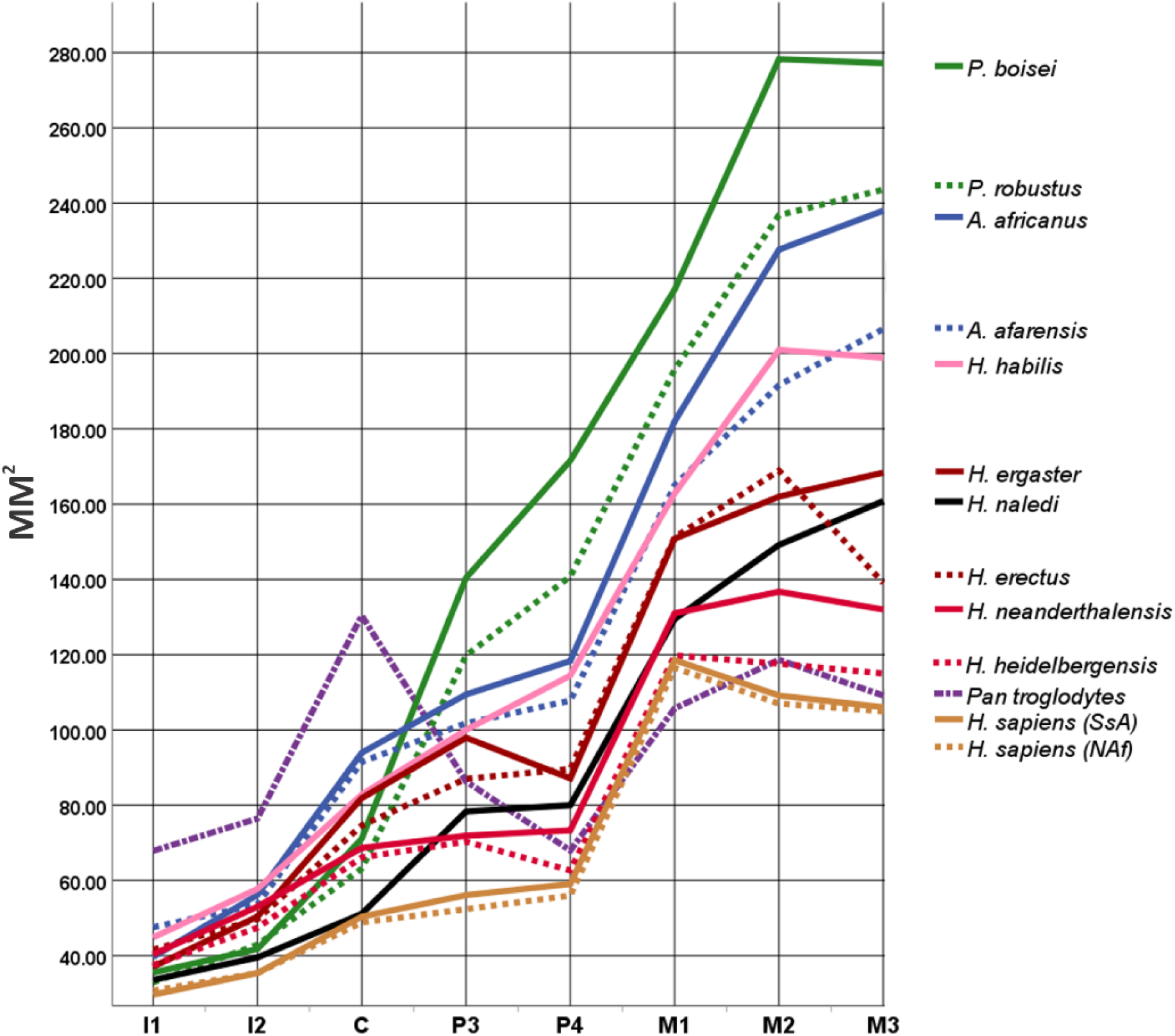
Line plot showing tooth-by-tooth trends in absolute occlusal surface areas of the mandibular dentition in mm^2^ by sample. Line colors and format apply loosely to genus (e.g., see Fig. 5, SI Fig. S1), but are primarily used to differentiate samples. See text for sample compositions.

The DM-corrected MD and BL measurements are listed in Table 2. When compared with Table 1, the effect of the scaling is clear. To illustrate, North African *H. sapiens* and *Pan* have the same mean UM1 MD diameter of 10.4 mm (Table 1), but their respective scaled values are 1.27 and 1.06 (Table 2); the latter value indicates a smaller UM1 in this dimension relative to all other teeth in the *Pan* dentition. On the other hand, North African *H. sapiens* and *P. boisei* have a corrected MD value of 1.38 for the LM1 (Table 2), yet the corresponding absolute dimensions are 11.2 and 15.5 mm (Table 2).

**Table 2.** DM_Raw size-corrected^a^ mesiodistal (MD) and buccolingual (BL) diameters for the samples used in the tooth size apportionment analyses.

|  | PAN<sup>b</sup> |  | PBO |  | PRO |  | AFA |  | AFR |  | HNA |  | HHA |  | HEG |  | HER |  | HHE |  | HNE |  | HSS |  | HSN |  |
| --- | --- | --- | --- | --- | --- | --- | --- | --- | --- | --- | --- | --- | --- | --- | --- | --- | --- | --- | --- | --- | --- | --- | --- | --- | --- | --- |
|  | MD | BL | MD | BL | MD | BL | MD | BL | MD | BL | MD | BL | MD | BL | MD | BL | MD | BL | MD | BL | MD | BL | MD | BL | MD | BL |
| UI1 | 1.20 | 0.96 | 0.82 | 0.62 | 0.83 | 0.68 | 1.01 | 0.79 | 0.96 | 0.77 | 1.03 | 0.71 | 1.05 | 0.71 | 1.12 | 0.77 | 1.03 | 0.84 | 1.02 | 0.83 | 1.03 | 0.90 | 1.04 | 0.85 | 1.05 | 0.86 |
| UI2 | 0.88 | 0.88 | 0.70 | 0.59 | 0.63 | 0.64 | 0.71 | 0.68 | 0.62 | 0.63 | 0.72 | 0.69 | 0.68 | 0.64 | 0.80 | 0.75 | 0.77 | 0.83 | 0.80 | 0.80 | 0.85 | 0.89 | 0.83 | 0.77 | 0.82 | 0.78 |
| UC | 1.13 | 1.27 | 0.77 | 0.78 | 0.79 | 0.86 | 0.94 | 1.02 | 0.92 | 0.95 | 0.89 | 0.94 | 0.92 | 0.97 | 0.96 | 0.96 | 0.96 | 1.04 | 0.95 | 1.04 | 0.93 | 1.07 | 0.92 | 1.00 | 0.92 | 0.99 |
| UP3 | 0.82 | 1.05 | 0.93 | 1.28 | 0.91 | 1.30 | 0.83 | 1.17 | 0.85 | 1.18 | 0.88 | 1.15 | 0.87 | 1.15 | 0.87 | 1.23 | 0.85 | 1.18 | 0.87 | 1.17 | 0.85 | 1.13 | 0.86 | 1.13 | 0.85 | 1.10 |
| UP4 | 0.73 | 1.03 | 1.03 | 1.45 | 0.98 | 1.41 | 0.86 | 1.17 | 0.85 | 1.24 | 0.89 | 1.21 | 0.88 | 1.15 | 0.84 | 1.17 | 0.81 | 1.17 | 0.85 | 1.18 | 0.83 | 1.13 | 0.81 | 1.12 | 0.82 | 1.12 |
| UM1 | 1.06 | 1.16 | 1.25 | 1.33 | 1.23 | 1.32 | 1.13 | 1.27 | 1.19 | 1.28 | 1.27 | 1.28 | 1.23 | 1.24 | 1.26 | 1.31 | 1.19 | 1.33 | 1.24 | 1.33 | 1.23 | 1.31 | 1.27 | 1.36 | 1.27 | 1.38 |
| UM2 | 1.04 | 1.18 | 1.38 | 1.50 | 1.32 | 1.47 | 1.22 | 1.38 | 1.29 | 1.44 | 1.34 | 1.40 | 1.23 | 1.37 | 1.25 | 1.32 | 1.17 | 1.34 | 1.22 | 1.39 | 1.16 | 1.33 | 1.20 | 1.37 | 1.19 | 1.36 |
| UM3 | 0.98 | 1.13 | 1.25 | 1.55 | 1.38 | 1.54 | 1.20 | 1.37 | 1.29 | 1.46 | 1.27 | 1.36 | 1.19 | 1.37 | 1.22 | 1.38 | 1.00 | 1.27 | 1.05 | 1.33 | 1.05 | 1.31 | 1.07 | 1.34 | 1.10 | 1.31 |
| LI1 | 0.79 | 0.89 | 0.47 | 0.60 | 0.51 | 0.57 | 0.63 | 0.67 | 0.56 | 0.60 | 0.67 | 0.59 | 0.61 | 0.65 | 0.56 | 0.64 | 0.66 | 0.65 | 0.62 | 0.70 | 0.59 | 0.76 | 0.64 | 0.69 | 0.65 | 0.71 |
| LI2 | 0.85 | 0.93 | 0.51 | 0.64 | 0.61 | 0.62 | 0.63 | 0.76 | 0.66 | 0.72 | 0.76 | 0.65 | 0.72 | 0.70 | 0.71 | 0.71 | 0.72 | 0.73 | 0.70 | 0.76 | 0.71 | 0.83 | 0.70 | 0.74 | 0.71 | 0.75 |
| LC | 1.08 | 1.25 | 0.70 | 0.80 | 0.72 | 0.77 | 0.83 | 0.98 | 0.85 | 0.93 | 0.78 | 0.78 | 0.85 | 0.87 | 0.91 | 0.88 | 0.84 | 0.93 | 0.82 | 0.92 | 0.83 | 0.93 | 0.84 | 0.89 | 0.81 | 0.90 |
| LP3 | 1.06 | 0.84 | 0.93 | 1.20 | 0.93 | 1.13 | 0.91 | 1.00 | 0.89 | 1.05 | 0.99 | 0.96 | 0.93 | 1.02 | 0.95 | 1.00 | 0.89 | 1.02 | 0.91 | 1.00 | 0.84 | 0.97 | 0.86 | 0.96 | 0.84 | 0.94 |
| LP4 | 0.80 | 0.88 | 1.15 | 1.18 | 1.03 | 1.20 | 0.93 | 1.04 | 0.94 | 1.06 | 0.95 | 1.00 | 0.99 | 1.06 | 0.87 | 0.97 | 0.89 | 1.05 | 0.83 | 0.99 | 0.83 | 1.00 | 0.87 | 1.00 | 0.86 | 0.99 |
| LM1 | 1.11 | 0.99 | 1.38 | 1.25 | 1.36 | 1.27 | 1.24 | 1.19 | 1.29 | 1.19 | 1.34 | 1.17 | 1.32 | 1.17 | 1.28 | 1.15 | 1.29 | 1.23 | 1.28 | 1.19 | 1.25 | 1.18 | 1.38 | 1.27 | 1.38 | 1.27 |
| LM2 | 1.16 | 1.07 | 1.56 | 1.41 | 1.50 | 1.39 | 1.35 | 1.27 | 1.44 | 1.33 | 1.46 | 1.23 | 1.48 | 1.30 | 1.34 | 1.19 | 1.36 | 1.30 | 1.31 | 1.23 | 1.28 | 1.20 | 1.30 | 1.24 | 1.32 | 1.23 |
| LM3 | 1.09 | 1.04 | 1.60 | 1.37 | 1.58 | 1.36 | 1.45 | 1.28 | 1.50 | 1.34 | 1.47 | 1.33 | 1.47 | 1.25 | 1.37 | 1.20 | 1.25 | 1.17 | 1.28 | 1.14 | 1.27 | 1.17 | 1.28 | 1.21 | 1.31 | 1.21 |

<sup>a</sup>See main text for details.
<sup>b</sup>PAN = *Pan troglodytes*, PBO = *P. boisei*, PRO = *P. robustus*, AFA = *A. afarensis*, AFR = *A. africanus*, HNA = *H. naledi*, HHA = *H. habilis*, HEG = *H. ergaster*, HER = *H. erectus*, HHE = *H. heidelbergensis*, HNE = *H. neanderthalensis*, HSS = *H. sapiens* (SsA), HSN = *H. sapiens* (NAf) (see text for details).

The effect of size correction is demonstrated further by submitting original (Table 1) and DM-values (Table 2) to separate UPGMA cluster analyses. With the former data, two large clusters are evident that, while indicating some likely links, e.g., *H. heidelbergensis* and *H. neanderthalensis*, and both *H. sapiens* samples, are based primarily on dentition size (Fig. 3). This determination is sustained by the crown area graphs (Figs. 1-2), where the ‘large’-toothed *P. boisei*, *P. robustus*, *A. africanus*, *A. afarensis*, *H. habilis*, and *H. ergaster* are on top, and ‘small’-toothed *H. naledi*, *H. erectus*, *H. heidelbergensis*, *H. neanderthalensis*, *Pan troglodytes*, and *H. sapiens* at the bottom. This arbitrary large-small dichotomy is quantified by summing all crown areas for each sample, where the former six range between 2483 and 1813 total mm^2^, and the latter six 1648 to 1154 mm^2^, as visualized in a bar graph (SI Fig. S1). In contrast, the UPGMA dendrogram of 32 DM-corrected values (Fig. 4) follows largely accepted phylogenies, as discussed below. *Pan* is separate from all other hominins, both *Paranthropus* species group together, and the other taxa group into two larger clusters; one contains the five most recent *Homo* samples (not including *H. naledi*), and the other cluster African-only species that, with one exception, date between ∼0.8 and 3.3 Ma. The exception is ∼335–236 ka *H. naledi*, in a cluster with *H. habilis* after size correction.

**Figure 3.**
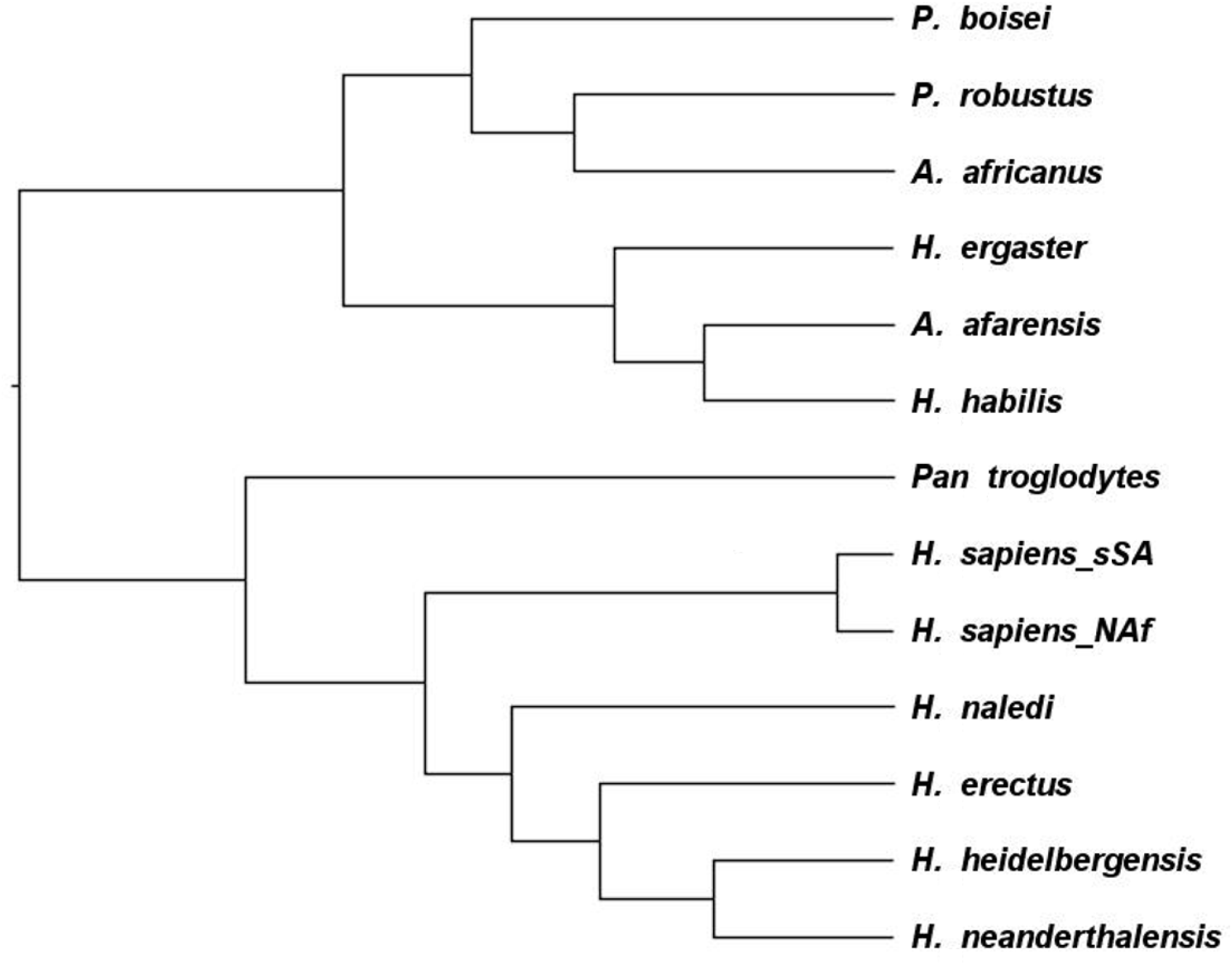
Average linkage (between groups) or UPGMA cluster analysis dendrogram based on 32 uncorrected MD and BL dimensions of the maxillary and mandibular teeth in *H. naledi* and 12 comparative samples. See text for methodological details.

**Figure 4.**
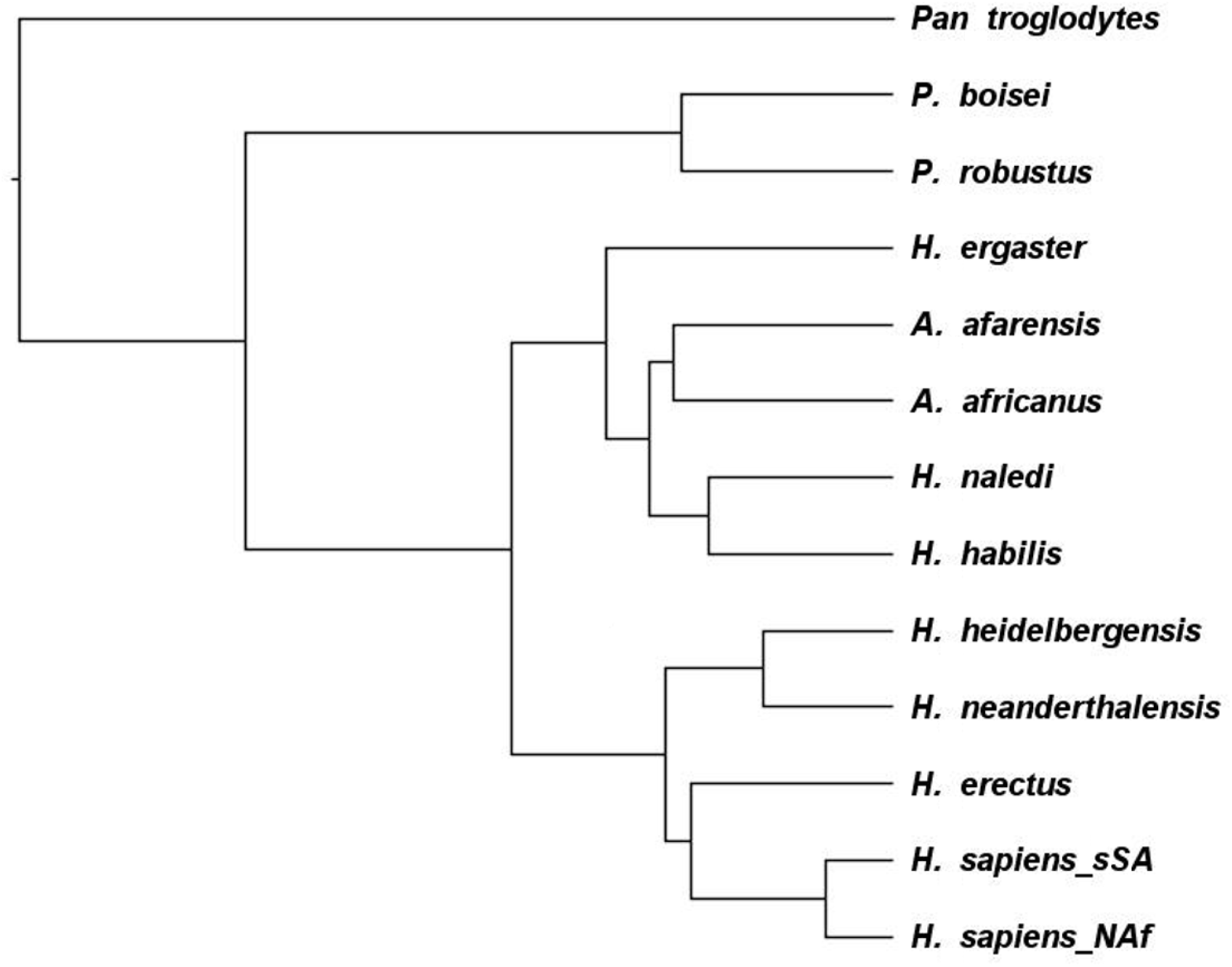
Average linkage (between groups) or UPGMA cluster analysis dendrogram based on 32 DM-corrected (i.e., scaled) MD and BL dimensions of the maxillary and mandibular teeth in *H. naledi* and the 12 comparative samples. See text for methodological details.

To complete the TSA analysis, the correlation matrix of DM_RAW-scaled data was submitted to PCA. The resulting un-rotated factor scores from three components having eigenvalues >1 were then used for plotting the among-sample variation in three dimensions. The component loadings, eigenvalues, individual variance, and total variance explained, i.e., 90.7%, are listed in Table 3. The loadings are also presented as bar graphs (SI Figs. S2-S4) to elucidate further those of most importance in driving the variation along each axis of the 3D plot (below). In other words, by interpreting this output one can determine how crown size is differentially apportioned or distributed along maxillary and mandibular tooth rows, to compare variation in patterning among species.

**Table 3.** Loadings^a^, eigenvalues, and the variance explained for the first three principal components based on size corrected maxillary and mandibular measurements in the 13 samples.

| Variable | Component |  |  |
| --- | --- | --- | --- |
|  | 1 | 2 | 3 |
| DM_MDUI1 | **-.916** | -.022 | -.310 |
| DM_MDUI2 | **-.786** | -.441 | .091 |
| DM_MDUC | **-.963** | .175 | .000 |
| DM_MDUP3 | **.831** | -.030 | .119 |
| DM_MDUP4 | **.933** | .142 | .156 |
| DM_MDUM1 | **.541** | **-.639** | -.412 |
| DM_MDUM2 | **.934** | .190 | -.240 |
| DM_MDUM3 | **.816** | .386 | -.251 |
| DM_BLUI1 | **-.923** | -.284 | .159 |
| DM_BLUI2 | **-.843** | -.391 | .191 |
| DM_BLUC | **-.978** | .060 | .131 |
| DM_BLUP3 | **.843** | .084 | .282 |
| DM_BLUP4 | **.923** | .160 | .272 |
| DM_BLUM1 | .483 | **-.852** | .011 |
| DM_BLUM2 | **.957** | -.085 | -.003 |
| DM_BLUM3 | **.969** | .077 | .039 |
| DM_MDLI1 | **-.873** | .136 | -.248 |
| DM_MDLI21 | **-.873** | .087 | -.390 |
| DM_MDLC | **-.886** | .344 | -.028 |
| DM_MDLP3 | -.217 | **.847** | -.183 |
| DM_MDLP4 | **.876** | .340 | .035 |
| DM_MDLM1 | **.766** | **-.500** | -.308 |
| DM_MDLM2 | **.953** | .196 | -.126 |
| DM_MDLM3 | **.945** | .238 | -.088 |
| DM_BLLI1 | **-.910** | -.033 | .254 |
| DM_BLLI2 | **-.928** | .011 | .283 |
| DM_BLLC | **-.873** | .300 | .312 |
| DM_BLLP3 | **.900** | .004 | .391 |
| DM_BLLP4 | **.923** | .075 | .293 |
| DM_BLLM1 | **.681** | **-.658** | .038 |
| DM_BLLM2 | **.927** | .027 | .176 |
| DM_BLLM3 | **.927** | .155 | -.172 |
| Eigenvalue | 23.756 | 3.703 | 1.577 |
| Variance (%) | 74.237 | 11.572 | 4.928 |
| Total Variance | 74.237 | 85.809 | 90.737 |

<sup>a</sup>Values in bold-face indicate strong loadings, i.e., $\geq |0.5|$ (see text for details).

The first component accounts for the greatest variance, 74.2%, identifying the main difference in relative, intraspecific tooth size. Like that of the first TSA hominin study (Irish et al., 2016), the familiar pattern of massive posterior and diminutive anterior teeth of both *Paranthropus* species was clearly identified. Thus, except for the DM-scaled BL dimension of the UM1, 0.483, and the scaled MD of the LP3, −0.217 (Table 3; SI Fig. S2), strong loadings (i.e., ≥|0.5|) of 0.541-0.969 indicate the relatively large cheek teeth, i.e., P3 to M3 in both isomeres; this influence pushed *P. boisei* and *P. robustus* toward the far positive end of the X-axis in Figure 5. The first observed exception—the DM-scaled BL dimension of UM1, along with the relatively moderate loading of 0.541 for the tooth’s scaled MD, marks the extreme M1<M2<M3 size progression in this genus. The second observed exception, the negative loading for the DM-scaled LP3 MD dimension, is more a function of the elongated sectorial premolar in *Pan* near the opposite end of the axis. Otherwise, the location of the latter species is driven by strong loadings of −0.786 to –0.978 for relatively large I1, I2, and C in both maxilla and mandible. The remaining 10 samples plot between these two extremes. The stimulus for this distribution is apparent in Table 2 where, except for UM1 and LM1, the two *Paranthropus* taxa have the largest size-corrected posterior diameters of all samples, especially compared to *Pan* with the smallest. The opposite is true for anterior teeth, where *Pan* has the largest scaled diameters, and *P. boisei* and *P. robustus* the smallest.

**Figure 5.**
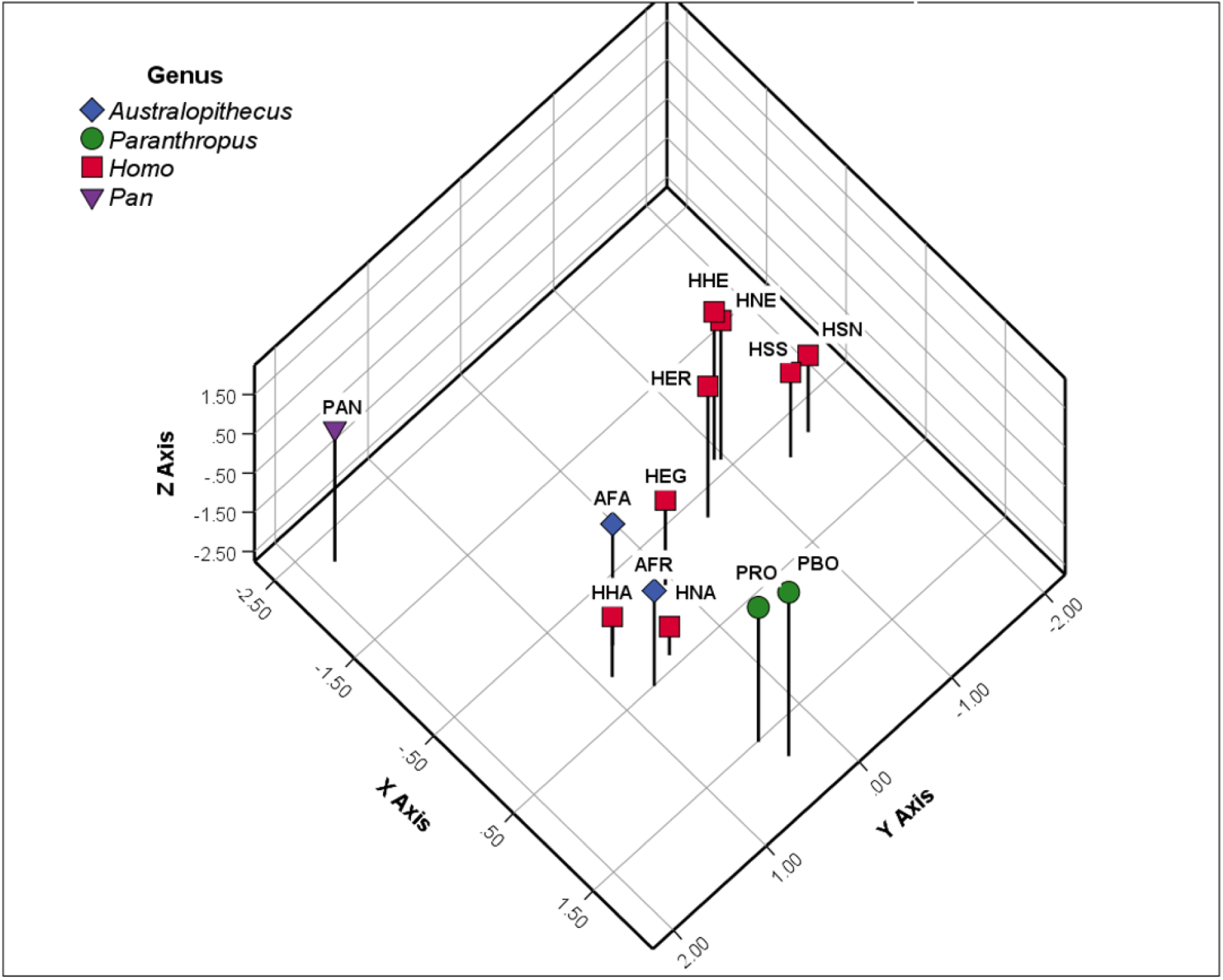
Three-dimensional ordination of retained principal component scores for tooth size apportionment (TSA) in the dentition of *H. naledi* (HNA) and 12 comparative samples. Accounts for 90.7% of the total variance (74.2% on X-axis, 11.6% on Y-axis, and 4.9% on Z-axis). AFA=*A. afarensis*, AFR=*A. africanus*, PRO=*P. robustus*, PBO=*P. boisei*, HHA=*H. habilis*, HEG=*H. ergaster*, HER=*H. erectus*, HHE=*H. heidelbergensis*, HNE=*H. neanderthalensis*, HSS=*H. sapiens* (sub-Saharan Africa), HSN=*H. sapiens* (North Africa), and PAN=*Pan troglodytes.* See text for methodological details and component descriptions.

The second component accounts for 11.6% of the total variance. The samples are separated by variation within the molar class. That is, as implied on component 1, maxillary and mandibular first molars are primarily responsible. As seen in Table 3 (and SI Fig. S3), the DM-scaled MD and BL values for these two teeth are both associated with strong negative loadings (−0.639, −0.852, −0.500, −0.658). Therefore, the M1s in both isomeres of the lowest scoring samples along the Y-axis are large relative to the corresponding M2s and M3s, to explain the location of most *Homo* species at the farthest, negative end (upper right of Fig. 5). In particular, *H. sapiens* has the characteristic M1>M2>M3 gradient, which is evidenced by the pertinent size-corrected MD and BL dimensions for UM1 and LM1 (see Table 2); this stands in contrast to samples near the positive end of the Y-axis, i.e., all australopith species. But obviously another factor is involved, viz. the strong loading of 0.847 for the DM-scaled MD of the LP3 (above); it drives *Pan* to the positive end of the axis, and affects *H. naledi* somewhat, with the latter’s slightly larger DM-scaled MD (0.99) than BL dimension (0.96) (Table 2). Again, these values are relative to those across the entire dentition, as seen by the less marked variation in actual MD (9.0 mm) and BL (8.8 mm) LP3 dimensions in *H. naledi* (Table 1). Though small sample size must be considered, also note the identical MD and BL dimensions for this tooth (10 mm) in *H. habilis* contra all remaining *Homo* species.

Finally, component 3 accounts for only 4.9% of the variance. There are no strong loadings, though several are of moderate magnitude (|0.3-.04|) (Table 3; SI Fig. S4). Thus, it is more difficult to interpret. However, low scoring samples on the Z-axis, such as *H. naledi* and *H. habilis*, are likely there in part because of: 1) larger DM-scaled MD (−0.310) relative to BL dimensions for the UI1, 2) larger DM-scaled MD (−0.412) relative to BL for the UM1, and larger DM-scaled MD (−0.390) relative to BL for LI2 than other species. That is, the three teeth may be characterized as relatively long and narrow. Near the top of the axis samples have a contrary pattern, while the DM-scaled MD dimension of LM1 (−.308) and the BL dimension of LP3 (0.391) are also involved relative to overall variation (refer to Table 2).

### 3.2 Bayesian phylogenetic inference with coded odontometric data

The cladogram from the strict-clock analysis is provided in Figure 6, with posterior parameters in Table 4. Like the dendrogram (Fig. 4) the *Paranthropus* species are sister taxa and two other hominin clades are again evident. The first clade contains the same five recent *Homo* samples, and the second, although polytomous, comprises the African-only species, including *H. naledi* and *H. habilis* as sister taxa. These relationships are noticeable in the Figure 5 scatterplot. Clade credibility values are 52.3-100%, indicating the proportion of trees in the MCMC sample with these clades. The lowest is for the node between *H. erectus* and the clade with sister taxa *H. heidelbergensis* and *H. neanderthalensis*. The highest value identifies four internal nodes between: 1) *Pan* and the hominins, 2) the two *Paranthropus* species, 3) the latter and all other hominins, and 4) both *H. sapiens* samples. As evident, the remaining values range from 55.7% to 97.8%, including 75.4% for *H. habilis* and *H. naledi*. Posterior probabilities diagnostics indicate a final tree number of 1.0, to provide support for a single tree (Ronquist et al., 2020).

**Figure 6.**
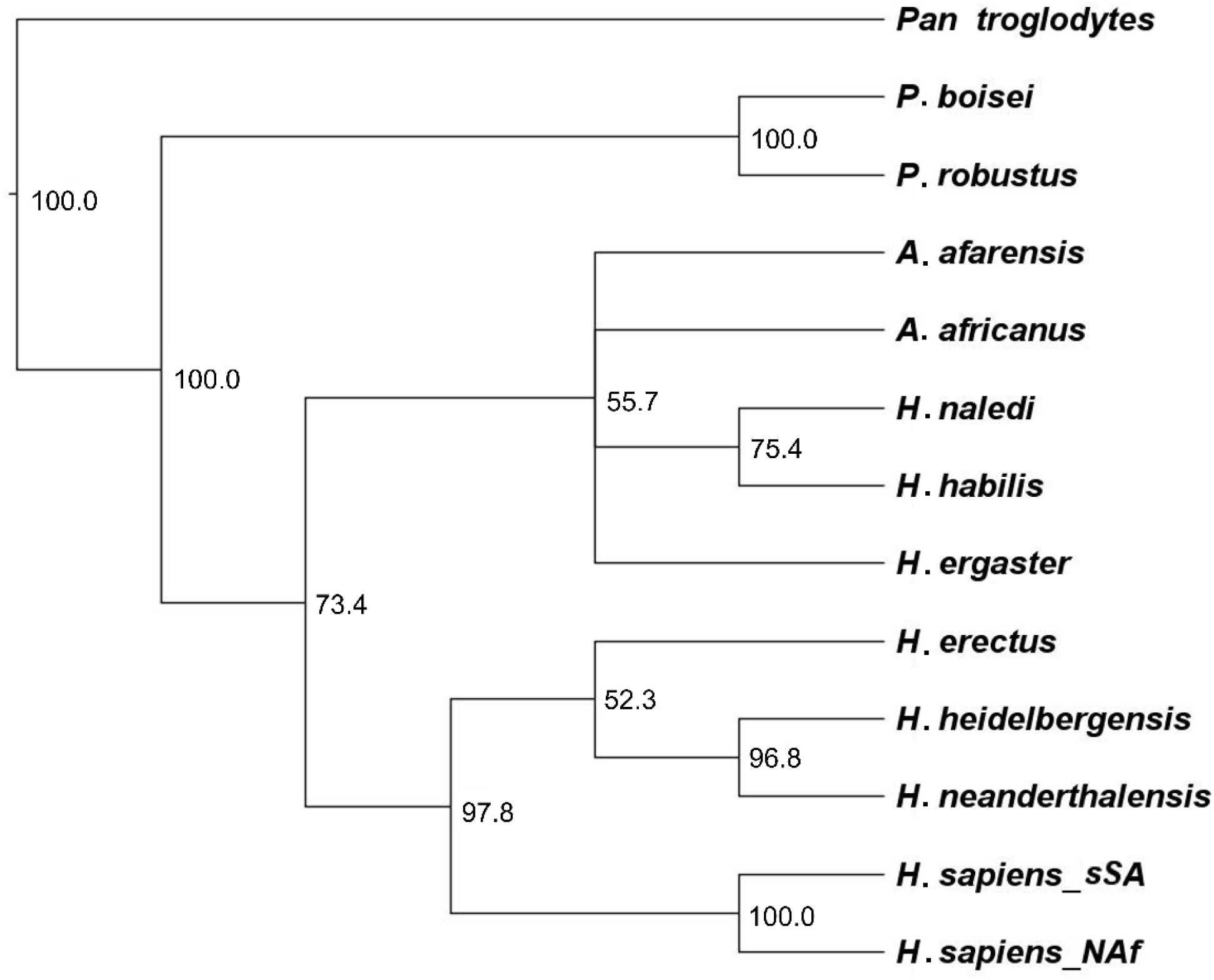
Bayesian inference cladogram from strict-clock analysis based on gap-weighted (i.e., coded), DM-scaled data under MKv, i.e., standard discrete model of *H. naledi* and 12 comparative samples, with clade credibility values for internal nodes included. See text for details.

**Table 4.**
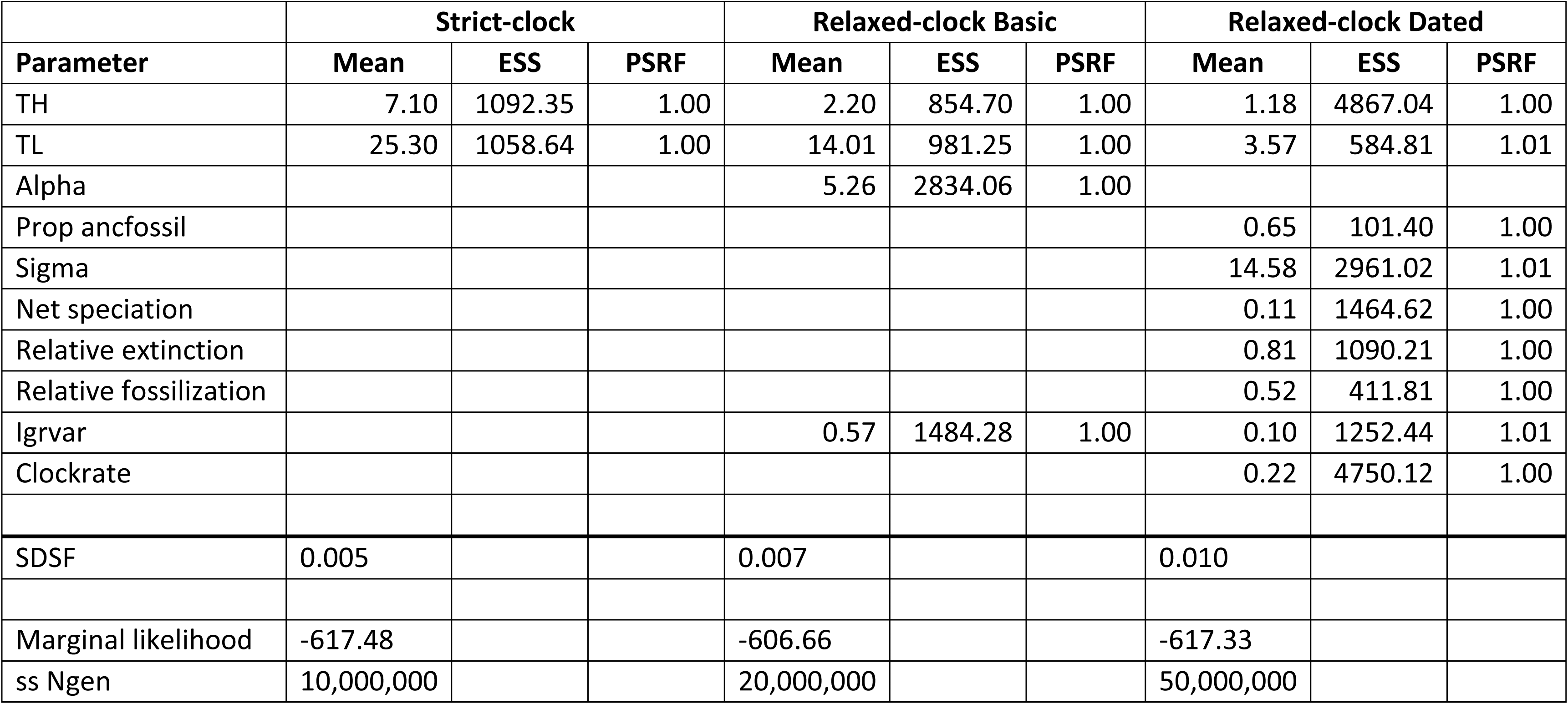
MrBayes posterior means, effective sample sizes (ESS), and potential scale reduction factors (PSRF), along with the average standard deviation of split frequencies (SDSF) and marginal log-likelihoods for determining the favored model.

|  | **Strict-clock** |  |  | **Relaxed-clock Basic** |  |  | **Relaxed-clock Dated** |  |  |
| --- | --- | --- | --- | --- | --- | --- | --- | --- | --- |
| **Parameter** | **Mean** | **ESS** | **PSRF** | **Mean** | **ESS** | **PSRF** | **Mean** | **ESS** | **PSRF** |
| TH | 7.10 | 1092.35 | 1.00 | 2.20 | 854.70 | 1.00 | 1.18 | 4867.04 | 1.00 |
| TL | 25.30 | 1058.64 | 1.00 | 14.01 | 981.25 | 1.00 | 3.57 | 584.81 | 1.01 |
| Alpha |  |  |  | 5.26 | 2834.06 | 1.00 |  |  |  |
| Prop anc fossil |  |  |  |  |  |  | 0.65 | 101.40 | 1.00 |
| Sigma |  |  |  |  |  |  | 14.58 | 2961.02 | 1.01 |
| Net speciation |  |  |  |  |  |  | 0.11 | 1464.62 | 1.00 |
| Relative extinction |  |  |  |  |  |  | 0.81 | 1090.21 | 1.00 |
| Relative fossilization |  |  |  |  |  |  | 0.52 | 411.81 | 1.00 |
| Igrvar |  |  |  | 0.57 | 1484.28 | 1.00 | 0.10 | 1252.44 | 1.01 |
| Clockrate |  |  |  |  |  |  | 0.22 | 4750.12 | 1.00 |
| SDSF | 0.005 |  |  | 0.007 |  |  | 0.010 |  |  |
| Marginal likelihood | -617.48 |  |  | -606.66 |  |  | -617.33 |  |  |
| ss Ngen | 10,000,000 |  |  | 20,000,000 |  |  | 50,000,000 |  |  |

An MCMC run of 2,000,000 generations for the basic relaxed-clock analysis was needed to achieve a standard deviation of split frequencies of <0.01 (Table 4); this value indicates that similar trees from each run and a representative sample from the posterior probability were obtained. As above, the maximum *a posteriori* tree is partially resolved, in this case with two polytomies. It also differs by position of the two *Paranthropus* samples, now nested within the African-only species clade (Fig. 7), and higher credibility values of 64.4-100%. The lowest identifies the node between *H. sapiens* and the other recent *Homo* samples. From there, it is 75.9% for the sister taxa *H. habilis* and *H. naledi,* 87.4% between the three early *Homo* and four australopiths from the African-only clade, and 87.9% for *H. heidelbergensis* and *H. neanderthalensis*. The other five node values are ∼100%.

**Figure 7.**
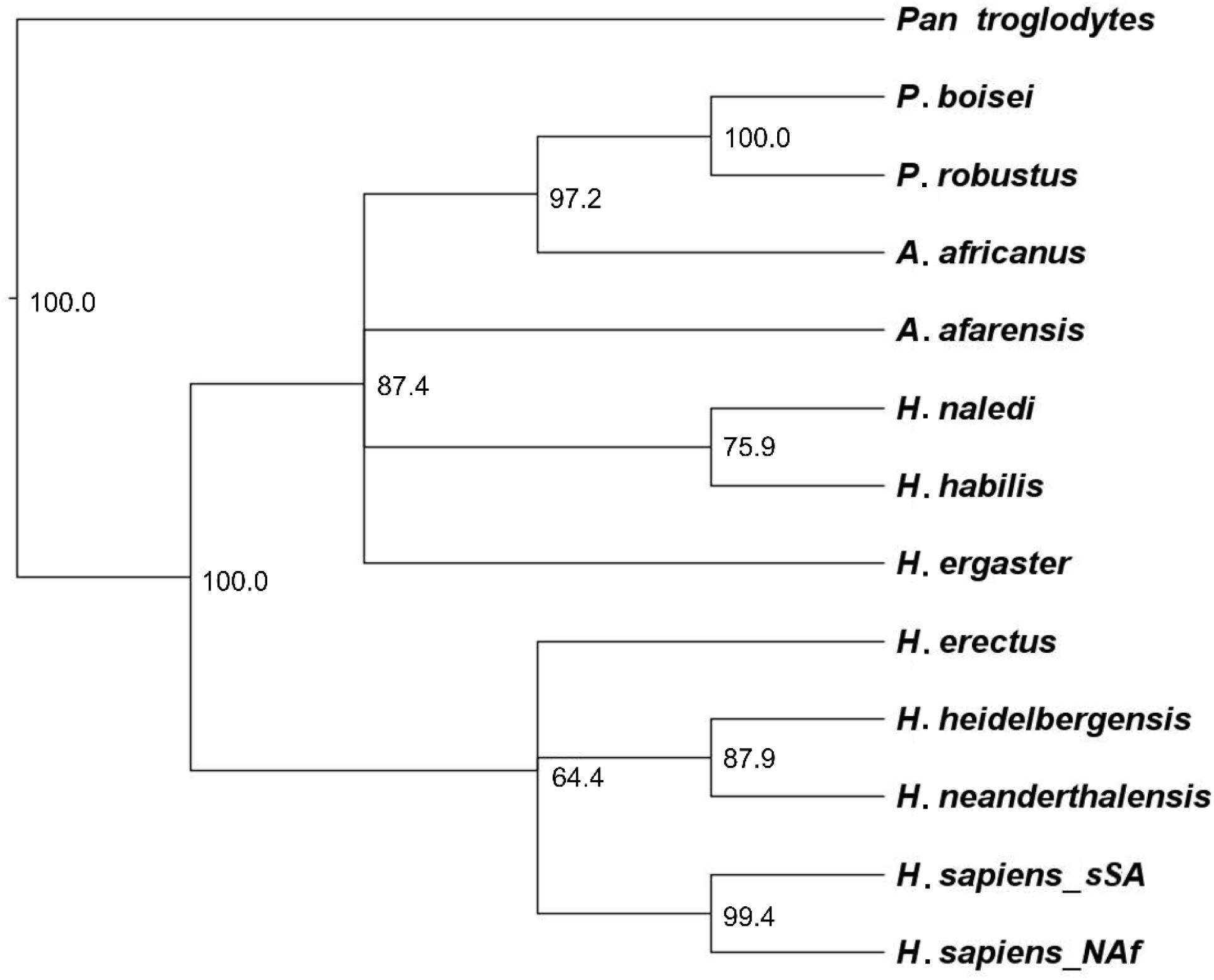
Bayesian inference cladogram from basic relaxed-clock analysis based on gap-weighted (coded), DM-scaled data under MKv model of *H. naledi* and the 12 comparative samples, with clade credibility values for internal nodes included. See text for details.

Finally, the dated relaxed-clock analysis using fossilized birth-death (FBD) branch lengths and lognormal rates priors produced the fully resolved phylogram in Figure 8 (also see Table 4), after a run of 5,000,000 generations. Rate variation is demonstrated by the branch lengths relative to origination and extinction dates. *Australopithecus afarensis* is an outgroup to the other hominins, while all *Homo* species form a clade separating them from the remaining australopiths—though with the lowest clade credibility of 54.7%. Other clade support values include 70.5% between *H. ergaster* and all later *Homo* species, to at or near maximal support for seven internal nodes, including *H. habilis* and *H. naledi* at 95.1%. The posterior probabilities diagnostics again provide support for a single tree.

**Figure 8.**
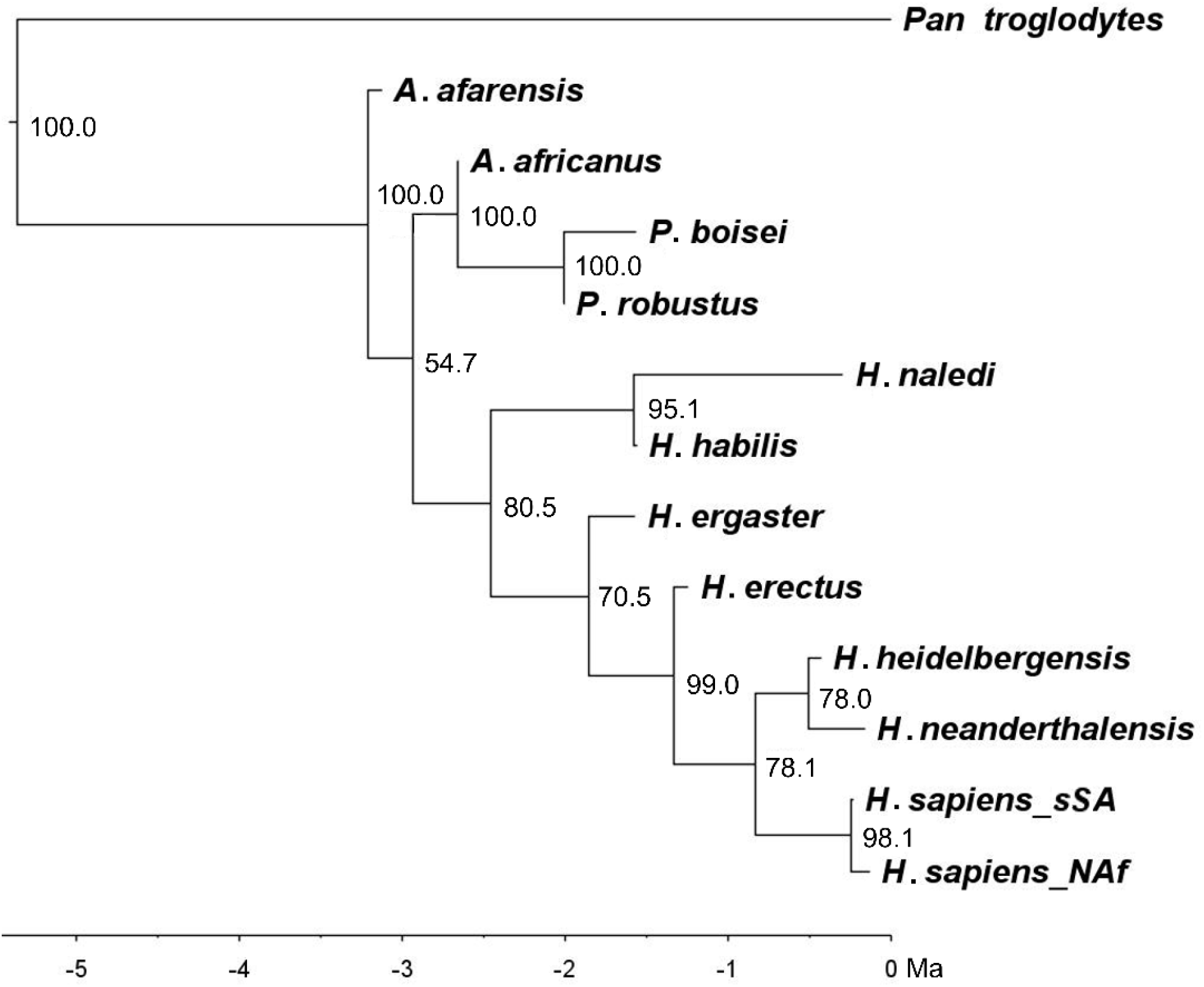
Bayesian inference phylogram from dated relaxed-clock analysis based on gap-weighted (coded), DM-scaled data under MKv model of *H. naledi* and the 12 comparative samples, with clade credibility values for internal nodes included. Scale in millions of years. See text for details.

The coded strict-clock, basic relaxed-clock, and dated relaxed-clock analyses yielded marginal likelihoods of −617.48 (M_1_), −606.66 (M_2_), and −617.42 (M_3_) for model comparison with Bayes factors (Table 4). Taking the difference between logarithms for each pair (M_1_ vs. M_2_, etc.) and doubling it [2*log*_e_*(B_12_)], the basic relaxed-clock model (M_2_) is favored. With criteria for ‘very strong’ evidence against or for a model a product of >10 (Kass and Raftery, 1995; Ronquist et al., 2020), those between the basic relaxed-clock and other coded models are 21.46 and 21.52, respectively. The product between the strict-clock model (M_1_) and the dated relaxed-clock model (M_3_) with lognormal rates, is 0.12, ‘not worth more than a bare mention’ evidence in favor of the latter.

### 3.3 Bayesian phylogenetic inference with continuous odontometric data

The strict-clock maximum *a posteriori* tree is presented in Figure 9, with RevBayes *a posteriori* diagnostics in Table 5. This fully resolved cladogram mirrors relationships resulting from the quantitatively coded data, showing two main hominin clades: sister taxa *P. boisei* and *P. robustus*, and all other hominins. Within the latter are two smaller clades, one with the five recent *Homo* species from the coded analyses, and the other containing African-only species, again with *H. habilis* and *H. naledi* as sister taxa. The clade credibilities range between 44.1% for the internal node linking *H. erectus* to the clade with *H. sapiens*, and 100% for nodes separating: 1) *Pan* from the hominins, 2) the *Paranthropus* species, 3) *H. heidelbergensis* and *H. neanderthalensis*, and 4) both *H. sapiens* samples. Node support of >90% was achieved in four other cases, including between the African-only and recent *Homo* clades.

**Figure 9.**
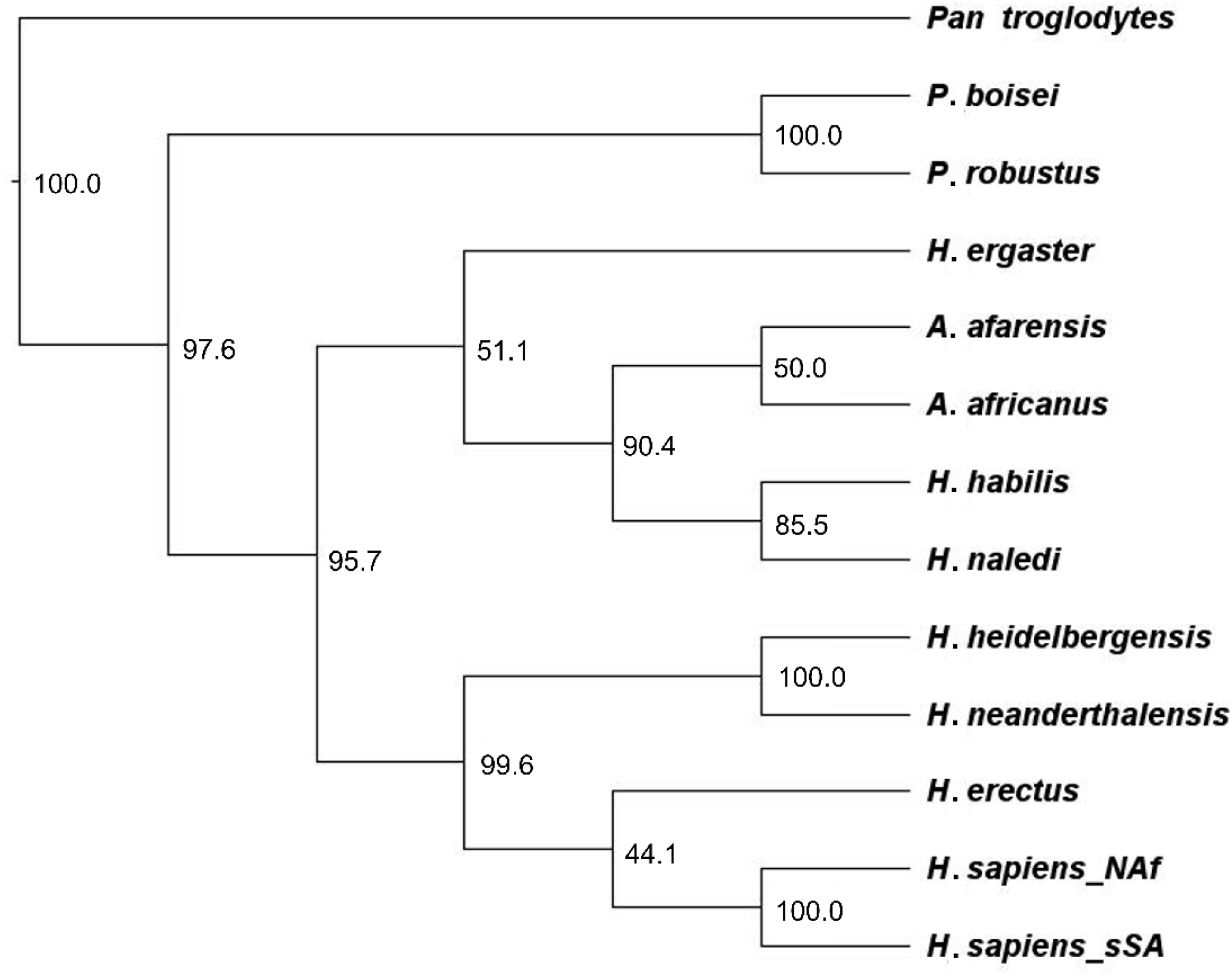
Bayesian inference cladogram from strict-clock analysis based on continuous DM-scaled data under Brownian-motion model of *H. naledi* and 12 comparative samples, with clade credibility values for internal nodes included. See text for details.

**Table 5.**
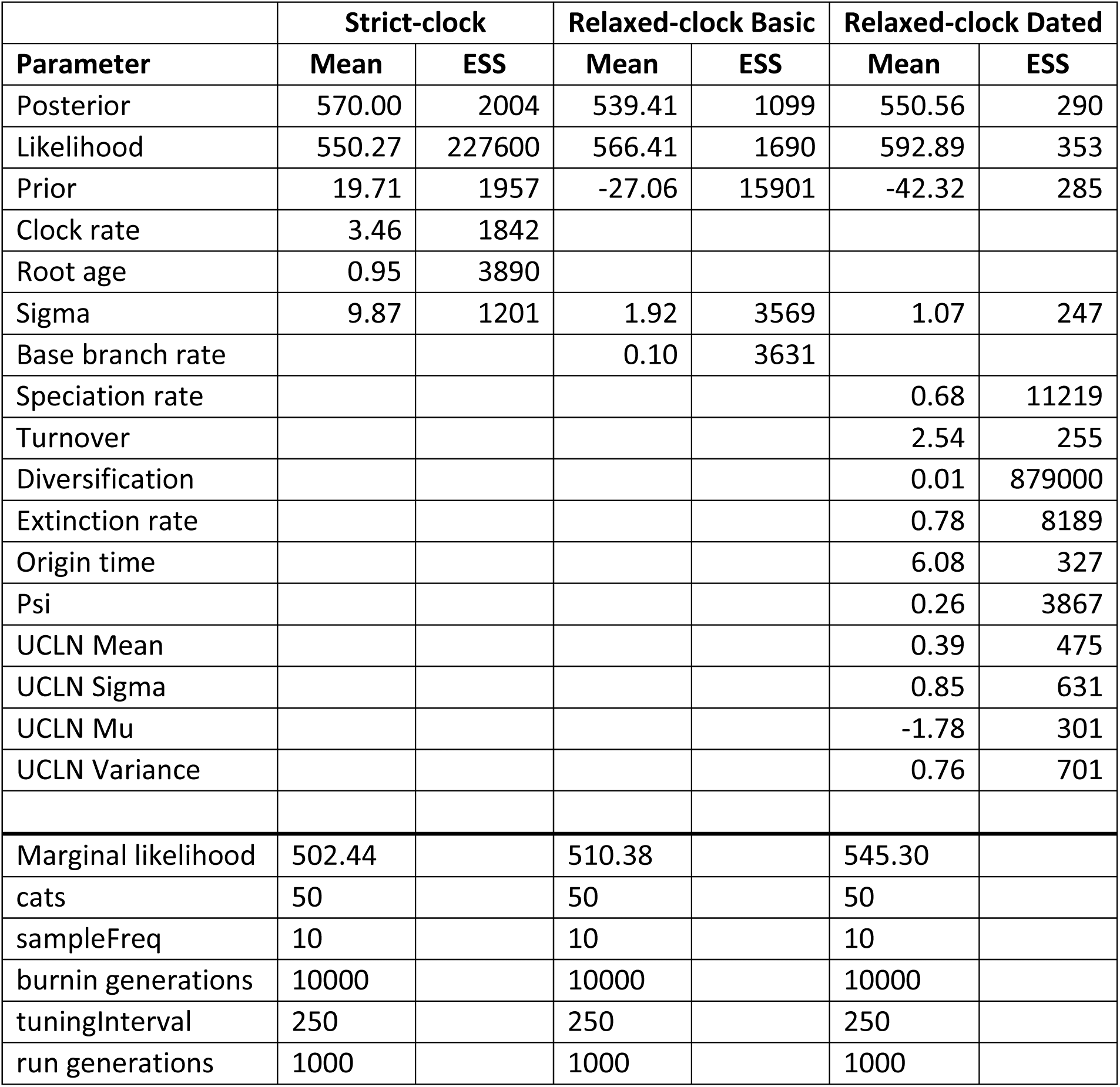
RevBayes parameter means and effective sample sizes (ESS), and those for the power posterior and stepping-stone analyses to calculate marginal log-likelihoods to determine the favored model.

The basic relaxed-clock *a posteriori* tree (Fig. 10) resembles that from the strict-clock. However, like the coded version of this phylogeny, the *Paranthropus* sister taxa are now nested within the African-only clade comprising *H. ergaster*, as an outgroup, followed by *A. afarensis,* and then *A. africanus, H. habilis*, and *H. naledi.* Clade support ranges from 48.5-100%, with the lowest at the node separating *A. africanus* from *H. habilis* and *H. naledi*. The credibility value between the latter two sister taxa is 68.6%. Near or maximal support is evident for other nodes between: 1) *Pan* and the hominins, 2) African-only and recent *Homo* species, 3) the two *H. sapiens* samples, 4) *H. erectus* and sister taxa *H. neanderthalensis* and *H. heidelbergensis*, 5) the latter two themselves, and 6) the two *Paranthropus* species.

**Figure 10.**
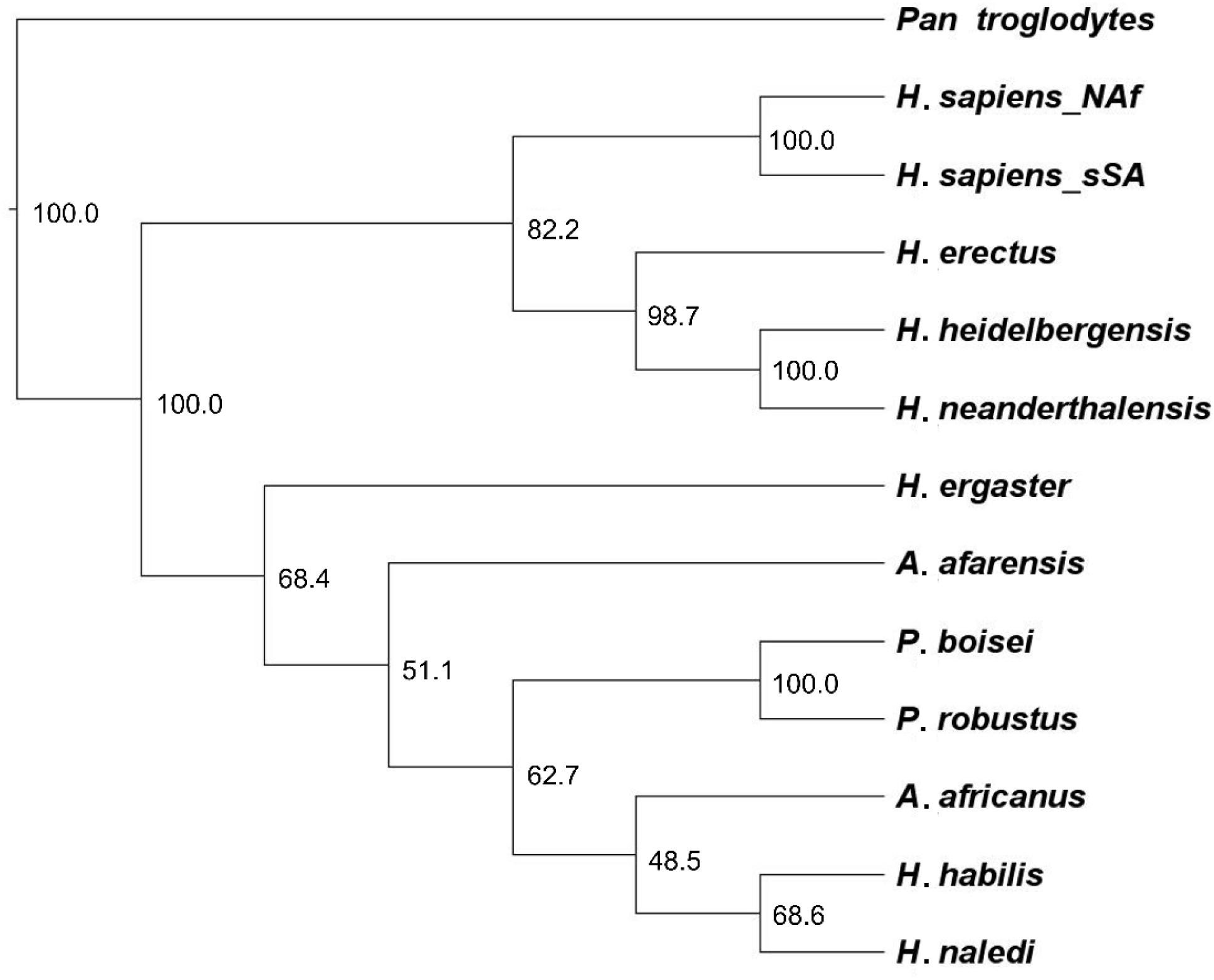
Bayesian inference cladogram from basic relaxed-clock analysis based on continuous DM-scaled data under Brownian-motion model of *H. naledi* and the 12 comparative samples, with clade credibility values for internal nodes included. See text for details.

Results of the relaxed clock fossilized birth-death model with dates (Fig. 11, Table 5) mirror those of the other continuous models. However, *A. afarensis* is the outgroup of all early African hominins and *H. naledi*, while *H. ergaster* is outgroup to the usual clade of five recent *Homo* samples, followed by *H. erectus*. Clade support is near maximal, with all nodes >92% except: 1) *H. ergaster* and the same five recent members of the genus (69.4%), 2) *H. erectus* and the four more recent *Homo* samples (76.6%), and 3) *H. heidelbergensis* and *H. neanderthalensis* clade from *H. sapiens* (64.4%). Rate variation is once more illustrated via branch lengths. As noted, to exclude the root from the hominin ingroup, *Pan* was forced to become ‘extinct’ prior to origination of the hominins using the 5-6 Ma age range.

**Figure 11.**
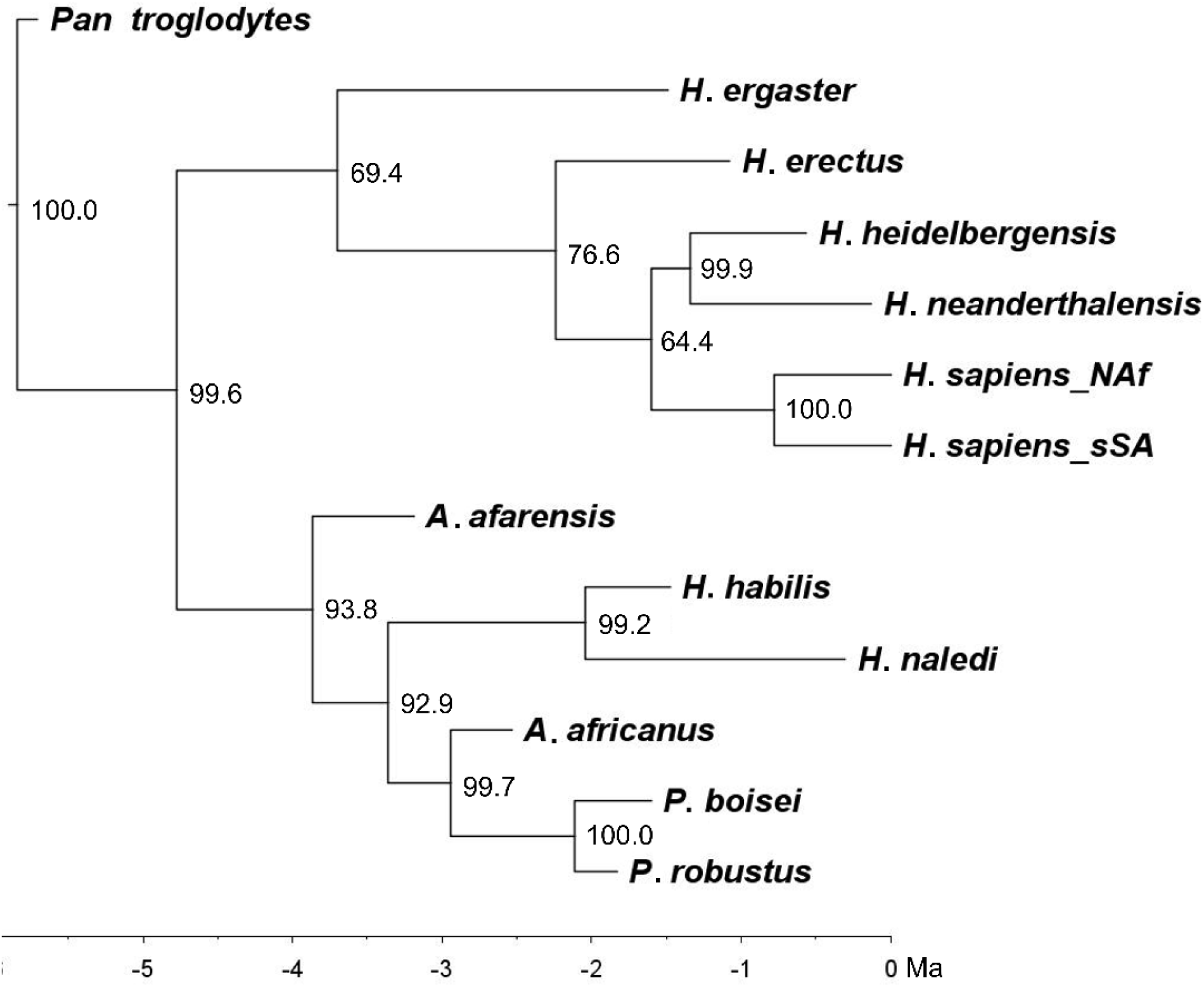
Bayesian inference phylogram from dated relaxed-clock analysis based on continuous DM-scaled data under Brownian-motion model of *H. naledi* and the 12 comparative samples, with clade credibility values for internal nodes included. Scale in millions of years. This is the preferred topology for this study. See text for details.

Finally, for the continuous strict-clock, basic relaxed-clock, and dated relaxed-clock analyses, marginal likelihoods of 502.44 (M_1_), 510.38 (M_2_), and 545.30 (M_3_) resulted for use in model comparison (Table 5). All log-likelihoods are positive, related to use of continuous characters (i.e., using probability densities vs. probabilities from discrete characters), though mainly due to DM-scaling of the odontometric data with small standard deviations (Gould et al., 2010; Canette, 2011; Cross Validated, 2020). For confirmation, and to assure improper priors (Baele et al., 2012) are not a factor, scaled data (Table 2) were replaced with mean dimensions from Table 1, and all stepping-stone analyses rerun. Corresponding marginal likelihoods from the non-scaled data are, as expected, negative, −543.94 (M_1_), −513.48 (M_2_), and −501.39 (M_3_), to support data as the root cause. Also as expected, both sets of marginal likelihoods are of analogous relative magnitude in characterizing the models, irrespective of data, i.e., M_3_>M_2_>M_1_. Focusing on the DM-scaled data, the difference between marginal likelihoods was again doubled to yield products for comparison: 15.88, favoring the basic relaxed-clock over the strict-clock model (M_2_>M_1_); 69.84, favoring the dated relaxed-clock over the basic relaxed-clock model (M_3_> M_2_); and 85.72, favoring the dated relaxed-clock over the strict-clock model (M_3_>M_1_). In all cases, evidence for M_3_ is ‘very strong’ (Kass and Raftery, 1995; Höhna et al., 2017a), with products ∼7X and 8.5X greater than the other analyses. In any case, the output is generally consistent among continuous runs, and with the coded results discussed above.

## 4. Discussion

The results address the three key points of this study mentioned at the outset: 1) the value of DM_RAW-scaled characters in assessing hominin relatedness, 2) the use of tooth size apportionment analysis and Bayesian inference with quantitatively coded and, particularly, continuous forms of the characters, and 3) insight from this extended analytical approach to understand better the evolutionary history of *H. naledi* relative to other fossil and recent species. Each point is discussed in turn.

### 4.1 The data

Qualitatively discretized craniodental data first published ∼30 years ago (Skelton et al., 1986; Skelton and McHenry, 1992) served as a basis for many subsequent studies on hominin relationships. This dataset was built on and modified (Strait et al., 1997; Strait and Grine, 2001) to yield, with additions (Collard and Wood, 2000), the Strait and Grine (2004) matrix. After amendments (Smith and Grine, 2008; Mongle et al., 2019) these, or similar qualitative craniodental data (e.g., Dembo et al., 2015, 2016), are the staples of modern hominin phylogenetic research. In the present study an entirely new approach is taken, using a set of continuous data (Table 2) derived from highly heritable crown dimensions—at least in modern humans, and largely alternative genetic pathways (Dassule et al., 2000; Shimizu et al., 2004; Sperber, 2006; Townsend et al., 2009; Harjunmaa et al., 2012). Though continuous, DM-scaling addresses the size component of crown dimensions; thus, undue influence on interspecific affinities (again see Figs. 3-4) or between the sexes is minimized. Instead, the patterning, or apportionment, of relative tooth size is identified among taxa for comparison. These dental characters—gap-weighted or continuous—hold several practical advantages over qualitative characters. Because teeth are relatively plentiful in the fossil record, MD and BL dimensions are readily available in the literature (as above). Further, based on long established protocol, these measurements are reasonably straightforward and the data collected comparatively unbiased among researchers. Observer replicability is a factor like all osteometric recording, but the subjective interpretation of morphological characters is avoided. Geometric morphometric and imaging techniques are also available to putatively increase the precision (Bernal, 2007; Kato and Ohno, 2009; Mitteroecker and Gunz, 2009; Gomez-Robles and Polly, 2012; Baab et al., 2012; Gomez-Robles et al., 2013; Braga, 2016; Hemphill, 2016a). That said, as used for the measurements in Table 1, calipers return comparable estimates of linear size and heritability of phenotype captured to higher-tech methods (e.g., Hlusko et al., 2002; Bernal, 2007).

Of course, no matter how advantageous a dataset might seem, it is only of value if the results are consistent, interspecific relationships plausible, and associated diagnostics supportive. Uniform cluster/clade membership is patent (Figs. 4, 6-11), with only minor differences among trees (discussed below). This consistency indicates that the results are data-driven, and not artifacts of the analytical method. The estimated relationships are also entirely credible, as evidenced by congruence with previous phenetic and cladistic studies for those species in common, *H. naledi* notwithstanding (below) (Strait et al., 1997; Strait and Grine, 2004; Smith and Grine, 2008; Berger et al., 2010; Irish et al., 2013, 2014, 2016, 2018; Dembo et al., 2015, 2016; Mongle et al., 2019). Beyond similarity, node support in the phylogenetic analyses is substantially stronger than earlier Bayesian studies (cf., Dembo et al., 2015, 2016; Mongle et al., 2019), and the posterior probabilities are indicative of a single maximum *a posteriori* tree in every case. According to Felsenstein (2004:299), “if the data strongly constrain the trees, then we might find only a few [or one] … accounting for most of the probability in the posterior,” vs. “if the data are fairly noisy, there might be millions of different trees.”

The consistency, plausibility, and strength of results from the DM-scaled data may be attributable to three factors. First, simply, there are no empty cells in the Nexus data matrix (e.g., see Dembo et al., 2016). Second, all morphological characters, even those qualitatively discretized, are “fundamentally quantitative,” and in this study they have been treated as such (Wiens, 2001:689; Felsenstein, 2004; Schols et al., 2004). Incomplete resolution affects two trees based on the six quantitatively coded states, but the continuous data afford a strong phylogenetic signal (Goloboff et al., 2006; Parins-Fukuchi, 2018a, 2018b). Third, while future research into how morphological characters from different regions affect inference is warranted (e.g., Kay, 2015; Dembo et al., 2016), focusing on one essential region arguably affords a clearer picture of evolution, to provide a more accurate phylogenetic signal than combining traits from across the skeleton.

Nevertheless, a case might be made that quantitative morphological characters are particularly susceptible to bias from evolutionary correlation—notably within an integrated structure, and homoplasy. Yet, as mentioned, they were said to be no more affected than qualitative discrete characters, and perhaps molecular data (Wiens, 2001; Parins-Fukuchi, 2018a, 2018b). Further, it is precisely because the data are recorded in serially homologous teeth, which function as a unit, that the scaled dimensions of each crown may actually be less subject to convergence, parallelism and, certainly, purported homoiology (Lycett and Collard, 2005; von Cramon-Taubadel, 2009). In any case, results matter, and dictate the usefulness of data, as expanded upon in the next section.

### 4.2 The analyses

As expected, the tooth size apportionment analysis results, both the preliminary cluster analysis (Fig. 4) of DM-scaled dimensions and submission of the latter to PCA with sample ordination (Fig. 5), emulate the prior study for those species in common (Irish et al., 2016). TSA analysis was literally made for continuous odontometric data, so the PCA results and 3D plot provide expedient phenetic estimates of relatedness. This output is equally valuable for phylogenetic analyses. That is, the component loadings (Table 3) can identify characters responsible for phenetic variation and, in Bayesian inference, clade formation. Scatterplots of the DM-scaled data by sample or, for direct pairwise comparisons, their quotients between samples (e.g., SI Fig. S7) can assist in visualizing interspecific character differences. As well, in conjunction with levels of cladogram node support, the 3D plot can discern those samples most likely to shift, i.e., so-called wildcard taxa (Nixon and Wheeler, 1992), or to remain constant. Unlike dendrograms and cladograms that emphasize the ‘end result’ of classification, sample ordination illustrates degrees of relationship, interpretable using a neighborhood approach (Guttman, 1954; Kruskal and Wish, 1978). For example, a likelihood for *H. ergaster* (HEG) to shift between the cluster/clade with the five recent *Homo* samples (excluding *H. naledi*), and that with African-only species (Figs. 4, 6-11), is apparent by its intermediate location between these large two groups in Figure 5. Similarly, the close proximity of four sample pairs in the figure (i.e., *P. boisei*/*P. robustus*, *H. naledi*/*H. habilis*, *H. heidelbergensis*/*H. neanderthalensis*, and both *H. sapiens* samples), indicates why all remain sister taxa across trees regardless of data type or model, with near maximal node support.

The three main hominin clades of the coded strict-clock tree (Fig. 6) are identical in composition to the three dendrogram clusters (Fig. 4), all with high clade credibility values. However, the cladogram has a polytomous node for daughter taxa *A. afarensis*, *A. africanus*, *H. ergaster*, and the *H. naledi*/*H. habilis* sister group. As well, *H. erectus* shifted slightly in position. The subsequent basic relaxed-clock model is favored over the strict-clock according to Bayes factors, and the cladogram (Fig. 7) indicates greater node support. However, two polytomous nodes are now evident, one at the base of the first major hominin clade of five recent *Homo* species, and the other at the second clade linking *A. afarensis*, *H. ergaster*, and the *H. naledi*/*H. habilis* sister taxa. Slight shifting of *H. erectus* and *A. africanus* occurred. The relationships are credible from both models, but lack of resolution and potentially unstable taxa indicate insufficient phylogenetic information (Maddison, 1989; Nixon and Wheeler, 1992; Purvis and Garland, 1993; Pol and Escapa, 2009). Though quantitatively coded, data compression likely remains a factor because of the six-state maximum when characters are ordered in MrBayes (Ronquist et al., 2020). Several trees in the studies using qualitatively coded data are likewise affected (Strait et al., 1997; Strait and Grine, 2004; Smith and Grine, 2008; Mongle et al., 2019). These issues, and another specific to gap-weighting are detailed below.

The final tree from coded DM-data, generated by the dated relaxed-clock model, is fully resolved with strong node support (Fig. 8). The position of *A. afarensis* is contrary to prior trees, but it still might be the most probable coded topology (also SI Fig. S5)—if not the favored model—assuming estimates from the studies with qualitatively discretized data are generally accepted. Excluding *H. naledi* in Dembo et al. 2016, the tree is fully congruent with those from prior Bayesian inference (Dembo et al., 2015; Mongle et al., 2019) and maximum parsimony (Strait et al., 1997; Strait and Grine, 2004; Smith and Grine, 2008; Mongle et al., 2019). For the model, any compromised signal from the limited number of states likely was improved by two factors. First, a relaxed-clock model is recommended for different species, as it can incorporate a prior distribution of evolutionary rates that vary among both taxa and branches (Felsenstein, 2004; Pybus, 2006). Second, the fossilized birth-death prior and dates promote unrestricted branch length variation. For example, a dated relaxed-clock analysis using default priors, most importantly equal rates and uniform branch lengths (not shown), produced a phylogram with clearly inaccurate branch variation relative to divergence times (SI Fig. S6) and, like the basic relaxed-clock tree it emulates (Fig. 7), two polytomies.

Moving to the Bayesian inference with continuous data, the strict-clock tree (Fig. 9), has two relatively low clade credibility values, but is completely resolved. It comprises the same three major clades as the coded version (Fig. 6), and matches the dendrogram (Fig. 4) exactly. That said the model, like the quantitatively coded variant, is least favored compared with the subsequent continuous models. Of these, the basic relaxed-clock tree is again fully resolved (Fig. 10), with the same two main hominin clades in Figure 7. Yet, *H. erectus* and *A. africanus* form different clades with the sister taxa *H. heidelbergensis*/*H. neanderthalensis*, and *H. habilis*/*H. naledi*, respectively. Further, the Bayes factor comparisons revealed strong evidence against it relative to the continuous dated fossilized birth-death model. Finally, the phylogram from the latter is fully resolved (Fig. 11), with exceptionally strong node support. The topology is basically consistent with previous trees. However, other than *A. afarensis* as outgroup to the African-only clade and shifting of *H. habilis*/*H. naledi* sister taxa, this tree is most like that from the coded dated model (compare Fig. 8).

Which phylogenetic estimate, then, is the most plausible? Based on the coded vs. continuous data, the answer should lie with Bayesian inference from the latter. Irrespective of practicality (i.e., data objectively collected, no need for ordered states), the trees (Figs. 6-11) support simulation studies that revealed continuous characters give lower topological reconstruction error than discrete traits, especially for morphological structures with high or very slow evolutionary rates (Parins-Fukuchi, 2018a; also see Wright and Hillis, 2014), and increasing amounts of trait correlation (Parins-Fukuchi, 2018a). So, while gap-weighting may be more objective than qualitative coding and all topologies are credible, at only six states it cannot accommodate the full range of continuous variation. To illustrate, *Pan troglodytes* (Table 2) has the smallest DM_RAW-scaled MD UP3 diameter (0.82) and *P. boisei* the largest (0.93), with a range of 0.12. The same diameter for the LM3 is again smallest in *Pan* (1.09) and largest in *P. boisei* (1.60), for a range of 0.51—nearly 5X greater. Yet for both characters, a state of 0 was assigned to *Pan*, and 5 to *P. boisei*. The magnitude in range is addressed across taxa but not the data, which can increase artificially the influence of characters with minimal variation, and vice versa. Thus, the partially resolved coded strict-clock and basic relaxed-clock trees may be discounted. The tree from the coded dated relaxed-clock model, though fully resolved and congruent with prior studies, was correspondingly shortchanged on phylogenetic signal.

Finally, regarding the continuous analyses, the strict-clock model is least favored and the separate *P. boisei*/*P. robustus* clade, though unsurprising according to Figure 5, is not seen in previous cladistic studies nor the continuous relaxed-clock analyses. An inadequacy of the strict-clock to accommodate taxa with dissimilar evolution rates (Felsenstein, 2004; Pybus, 2006), which like phenetic analyses serves to focus the comparisons on general similarity, may play a role. The continuous basic relaxed-clock model is favored over the strict-clock, but some topological issues were noted. That leaves the continuous dated relaxed-clock results. Bayes factor comparisons indicate the strongest evidence for this model, its tree is again fully resolved, and node support is highest. As such, this topology (Fig. 11), in reference to all others, is deemed most plausible for assessing the phylogeny of *H. naledi* and the comparative taxa.

### 4.3 *The phylogeny of* Homo naledi

The general congruence of hominin relationships with past studies has been covered thoroughly, so the emphasis here is on *H. naledi.* It seems that a common supposition (e.g., Greshkho, 2017), yet with minimal published support, is that the species is a descendant of African *H. erectus* (*H. ergaster*). However, in the original paper Berger et al. (2015) describe only what was considered to be enough similarities with several *Homo* species, including *H. erectus,* to warrant classification in the genus. Using published craniometric data Thackeray (2015) agreed, although he found *H. naledi* most similar to *H. habilis*, and to a lesser extent *H. rudolfensis* and *H. ergaster*. Overall, the comparative analyses of crania and post-crania indicate *H. naledi* exhibits both *Homo*- and *Australopithecus*-like features. Examples include a well-developed, arched supraorbital torus separated from the vault by a continuous supra-toral sulcus like in *H. habilis* and *H. erectus*, marked angular and occipital tori like *H. erectus*, and facial similarities to *H. rudolfensis* (Berger et al., 2015; Hawks et al., 2017; Schroeder et al., 2017). The cranium is nothing like recent *Homo* species—as indicated by its endocranial morphology (Holloway et al., 2018) and small cranial capacity like *Australopithecus* (Garvin et al., 2017). In the post-crania, *Homo*-like features include long tibiae and gracile fibulae, muscle attachments suggestive of a striding gate, and modern traits in the ankles, feet, and hands. *Australopithecus*-like traits include curved phalanges, a wide lower thorax, ape-like arms, primitive pelvic morphology, and the same concerning certain aspects of the femora (Berger et al., 2015; Harcourt-Smith et al., 2015; Kivell et al, 2015; Feuerriegel et al., 2017; Garvin et al., 2017; Hawks et al., 2017; Marchi, 2017; Williams et al., 2017; VanSickle et al., 2018).

This mosaic of plesiomorphic, apomorphic, and apparent autapomorphic characters affected the prior attempt at Bayesian inference by Dembo et al. (2016). In their phylogram *H. naledi* is nested within a clade of 11 *Homo* species and *A. sediba*, but its position therein is ambiguous. The species cannot be excluded as a sister taxon to any one of several clade members, including *H. antecessor*, *H. erectus*/*ergaster*, *H. habilis*, *H. floresiensis*, and *H. sapiens*, among others. Node support between *H. naledi* and a smaller clade containing *H. antecessor*, *H. sapiens*, and the sister taxa *H. heidelbergensis* and *H. neanderthalensis*, is only 36%. Other clade credibility values leading to the latter node range between 21-54% (Dembo et al., 2016). A phenetic comparison of dental morphological traits also found *H. naledi* to group nearest *H. habilis* and *H. ergaster*. However, the unique combinations and expressions of traits differ enough to support its taxonomic status as a separate species in the genus (Irish et al., 2018). As well, the species’ molar root metrics revealed similarities with individual specimens of *H. habilis* (KNM-ER 1805), *H. ergaster* (SK 15), and early *Homo* sp. (SK 45) (Kupczik et al., 2019).

Most recently, research into dental morphological similarities with other hominins has tacked toward *Paranthropus*. For example, the deciduous lower canine and upper and lower first molars of *H. naledi* share apparent derived traits with the latter genus, though features of the second deciduous molars are *Homo*-like (Bailey et al., 2019). In a geometric morphometric study of lower premolar enamel-dentine junctions (EDJ), Davies et al. (2020) report the species is closest to *P. robustus* in a PCA ordination of the first two components (73.7% of total variation) for LP3 shape. *Homo habilis* is plotted nearby, but other specimens in the genus, including *H. erectus* and, in particular, more recent *H. neanderthalensis* and *H. sapiens*, are increasingly distinct. That said, the third component (6.4% of variation) of LP3 shape separates *H. naledi* and *P. robustus*, as do analyses of LP4 EDJ morphology and the centroid sizes of both premolars. From this, the authors maintain that the suite of traits is distinguishable from that of other hominin specimens in their analysis, including most early and later *Homo* species. Again, the mosaic nature of *H. naledi* is evident.

*Homo naledi* is seemingly a morphometric enigma. Cranial, dental, and post-cranial features offer conflicting evidence for the species’ exact place in hominin evolution—though with enough agreement to assign it to genus. Here, the DM-scaled MD and BL data indicate the species is firmly linked to *H. habilis* though, again, small sample size must be considered. Despite the method or data, i.e., quantitatively coded or continuous, the two remain sister taxa with high node support of ∼70-99%; the latter value is from the preferred continuous dated relaxed-clock phylogram. Further, with exception (Fig. 8), *H. naledi* is always nested in the cluster/clade of the much older African species *A. afarensis*, *A. africanus*, and *H. habilis* (Figs. 4, 6-7, 9-11), which also often includes *H. ergaster* (Figs. 4, 6-10), as well as *P. robustus* and *P. boisei* in some trees (Figs. 7, 10-11). Other than Figure 8, with *H. naledi* and its sister taxon as an outgroup, the species does not form a clade with *H. erectus*, *H. heidelbergensis*, *H. neanderthalensis*, and *H. sapiens*.

Looking at how size is apportioned along the dental rows, *H. naledi* and other *Homo and Australopithecus* species are characterized by general uniformity compared to extreme opposing patterns in *Pan troglodytes* and *Paranthropus* (Tables 2-3, Fig. 5 X-axis). Contra *Pan*, *H. naledi* has smaller anterior and larger posterior teeth. On an individual basis, other than *Pan*’s sectorial LP3, the teeth of *H. naledi* also have relatively larger DM-scaled MD than BL diameters (SI Fig. S7a). A pattern contrary to highly derived *P. robustus* and *P. boisei* would then be expected in *H. naledi*, i.e., larger anterior and smaller posterior teeth. This pattern is evident, but enough of the scaled dimensions are similar to *Paranthropus*, most notably *P. robustus* (Table 2), that exceptions occur. That is, UM1s are equivalent in relative size across these species, as are DM-scaled MD dimensions of the UP3, UM2, LM1, and LM2, and DM-scaled BL equivalents of the UI1, LI1, LC, and LM3 (SI Fig. S7b-c). Again, as with *Pan*, *H. naledi* teeth are comparatively longer in DM-scaled MD dimensions than, in this instance, the buccolingually expanded teeth of *Paranthropus*.

The apportionment of tooth size in *H. naledi* is most similar to that of *H. habilis* and, to a lesser degree, *A. africanus* and *A. afarensis.* Other than the general uniformity in DM-scaled anterior and posterior dentition size, all four have a strong M1<M2<M3 gradient in contrast to the diachronic trend toward M1>M2>M3 that typifies modern humans (Tables 2-3, Fig. 5 Y-axis). As well, the molars and a number of other teeth are of similar relative size among these four species. However, some scaled dimensions of individual teeth distinguish australopithecines from *H. naledi*. The latter has a noticeably smaller LC, yet comparatively large scaled MD dimensions of the UI2, LI1, and LI2—particularly in contrast to *A. africanus* (SI Fig. S7d-f). Though less marked than in *Pan* (above), the scaled MD dimension of the *H. naledi* LP3 is also large vs. the BL, as indicated by the associated high loading in Table 3. *Homo naledi* can be differentiated from *H. habilis* on these bases to some extent, but their similarities are more apparent. As indicated, they are the only two hominin species with an LP3 that is not wider (BL) than it is long (MD) (Table 1). In fact, the teeth in both species are characterized by large DM-scaled MD dimensions relative to all australopiths (Tables 2-3, Fig. 5 Y- and Z-axes). Beyond the shared molar size gradient, *H. naledi* and *H. habilis* also have long and narrow posterior teeth (SI Fig. S7f) in comparison to derived recent *Homo*, which exhibit mesiodistally reduced premolars and molars (SI Fig. S7g-l). However, anterior teeth of the latter species are relatively larger in both isomeres, particularly BL dimensions, than *H. naledi* or *H. habilis*.

Based on these characters, which serve to link *H. naledi* to the most ancient hominin species included in the analysis (Fig. 11) [the potential new date for African *H. erectus* not withstanding (Herries et al., 2020)], *H. naledi* exhibits an apparent plesiomorphic pattern of size apportionment. This inference, of course, assumes ancestry of *A. afarensis* and/or *A. africanus* to *H. habilis*, which in turn is representative of the basal member in its assigned genus. Other researchers have recently made similar statements on the basis of alternate skeletal structures. Schroeder et al. (2017) report that while certain cranial features ally *H. naledi* with *H. erectus*, those of the mandible are closer to basal *Homo.* Similarly, Holloway et al. (2018:5741) note that “derived aspects of endocranial morphology in *H. naledi* were likely present in the common ancestor of the genus.” And, Davies et al. (2020) suggest that several derived features of the premolars shared by *H. naledi* and [African] *H. erectus*, are actually homoplastic, evolving independently in these species from a basal *Homo* condition like *H. habilis*. They conclude by proposing “*H. naledi r*epresents a long surviving lineage that split from other members of the genus *Homo* relatively early” (Davies et al., 2020:13196.9). The present results support this inference and others finding links to a common, and early, *Homo* condition, based on the use of DM_RAW-scaled odontometric data in phenetic and, for the first time in fossil hominin research, Bayesian inference of continuous characters. The phylogenetic place of *H. naledi* is becoming clearer. More remains are being recovered and analyzed, but of greatest importance for increasing clarity is the discovery of specimens older than the age of the Dinaledi chamber; as implied by the above findings, they should be present somewhere in South Africa, if not farther afield. As/if more ancient, reliably dated *H. naledi* remains are obtained, it will be possible to discern just how long this putative long surviving lineage survived, either alongside or in the shadow of multiple successive hominin species, including *H. sapiens*.

## 5. Summary

The DM_RAW mathematical correction suggested in Jungers et al. (1995) was used to equivalently scale 32 standard MD and BL crown measurements in *H. naledi* relative to 12 other Plio-Pleistocene and recent samples. The aim was to provide further characterization of the recently discovered South African hominin. The comparative species are: *A. africanus*, *A. afarensis*, *P. robustus*, *P. boisei*, *H. habilis*, *H. ergaster*, *H. erectus*, *H. heidelbergensis*, *H. neanderthalensis*, two samples of *H. sapiens*, and *Pan troglodytes*. The DM-scaled data were employed in UPGMA cluster analysis and an approach called tooth size apportionment (TSA) analysis to assess inter-sample phenetic affinities (Irish et al., 2016). Then, for the first time, these quantitative characters were used with Bayesian inference. Phylogenetic relationships were initially estimated based on quantitatively-coded (gap-weighted) versions of the scaled data under an Mkv model. In another first concerning hominin analyses, phylogenies were inferred based on the continuous data themselves in Bayesian inference under a Brownian-motion model. The latter approach proved to be superior, but the range of output, whether dendrogram, 3D ordination, or phylogenetic tree—irrespective of Bayesian priors, returned highly comparable results. Relationships among comparative samples are congruent with those in earlier phylogenetic studies based on discrete characters, though with greater node support. With regard to *H. naledi*, the species forms a clade with *H. habilis* as sister taxa. With exception, it is always nested within the cluster/clade of much older African species that, based on published dates, range between 3.3 Ma to ∼800 ka. These are: *A. afarensis*, *A. africanus*, and *H. habilis*, along with *H. ergaster* in many trees, as well as *P. robustus* and *P. boisei* to a lesser extent. Other than the noted exception, *H. naledi* (as well as its sister taxon, *H. habilis*) does not form a clade with the more recent *H. erectus*, *H. heidelbergensis*, *H. neanderthalensis*, and *H. sapiens* species, in agreement with several recent studies based on comparisons of alternative skeletal features. *Homo naledi* has a plesiomorphic pattern of tooth size apportionment, like the most ancient species in the study and contra that of more recent and living members of its genus. This apparent basal *Homo* condition implies that the origin of *H. naledi* predates the ∼335–236 ka age of the Dinaledi Chamber from which the original fossils were recovered (Dirks et al., 2017). The species may indeed represent a long-lived side branch in the genus *Homo*, perhaps rivaling *H. habilis* or another basal species in age, while persisting until the advent of *H. sapiens*.

## Acknowledgements

Thank you to the individuals currently or formerly affiliated with the institutions at which the odontometric measurements of North and sub-Saharan *H. sapiens* samples were recorded. These include: Charles Merbs and Donald Morris from Arizona State University (ASU); Elden Johnson from the University of Minnesota; Douglas Ubelaker, David Hunt, and Carol Butler from the National Museum of Natural History; Ian Tattersall, Jaymie Brauer, and Gary Sawyer from the American Museum of Natural History; Andre Langaney, Frances Roville-Sausse, and Miya Awazu Periera da Silva from the Musée de l’Homme; Romuald Schild, Michal Kobusiewicz, and Jacek Kabaciński from the Combined Prehistoric Expedition to Gebel Ramlah; and Renée Friedman from the Hierakonpolis Expedition. Thanks are also extended to Lee Berger, from the Evolutionary Studies Institute and Centre for Excellence in PalaeoSciences; Darryl de Ruiter, Texas A&M, for those measurements presented in our 2016 TSA publication that are incorporated here; and Lucas Delezene, University of Arkansas, who provided a list of publications containing Asian *H. erectus* MD and BL measurements, several of which were accessed for the present summary data. Funding for the data collection by the first author came from the National Science Foundation (BNS-9013942), the ASU Research Development Program, American Museum of Natural History, Friends of Nekhen, and a National Science Foundation grant awarded to Jerome Rose (BCS-0119754).

