## Supplementary Information for "Tooth size apportionment, Bayesian inference, and the phylogeny of *Homo naledi*"

**SI Table S1**

MrBayes parameters and settings/priors used for the three Bayesian inference clock analyses based on the quantitatively-coded DM-scaled data.

| | **Strict-clock** | **Relaxed-clock Basic** | **Relaxed-clock Dated** |
|---|---|---|---|
| **Parameter**[a] | | | |
| Ngen | 1,000,000 | 2,000,000 | 5,000,000 |
| Rates | Equal | Gamma | Log-normal |
| Nlnormcat | | | 4 |
| Sigma | | | Exponential (1.0) |
| Coding | Variable | Variable | Variable |
| Ctype | Ordered | Ordered | Ordered |
| Statefreqpr | Dirichlet | Dirichlet | Dirichlet |
| Symdirihyperpr | Fixed(Infinity) | Fixed(Infinity) | Fixed(Infinity) |
| Topologypr | Uniform | Constraints(hominins) | Constraints(Pan,hominins) |
| Brlenspr | Clock:Uniform | Clock:Uniform | Clock:fossilization |
| Treeagepr | Gamma(1.0,1.0) | Gamma(1.00,1.00) | |
| Speciationpr | | | Exponential(10.0) |
| Extinctionpr | | | Beta(1.0,1.0) |
| Fossilizationpr | | | Beta(1.0,1.0) |
| SampleStrat | | | Random |
| Sampleprob | | | 1 |
| Nodeagepr | Unconstrained | Unconstrained | Calibrated |
| Clockratepr | Fixed(1.0) | Fixed(1.0) | Normal(0.20,0.02) |
| Clockvarpr | Strict | Igr | Igr |
| Igrvarpr | | Exponential(10.0) | Exponential(10.0) |

[a]Any parameters not specifically mentioned in the text use default MrBayes values (e.g., flat priors)

**SI Table S2**

RevBayes parameters and settings/priors used for the three Bayesian inference clock analyses based on the continuous DM-scaled data.

| | **Strict-clock** | **Relaxed-clock Basic** | **Relaxed-clock Dated** |
|---|---|---|---|
| **Parameter**[a] | | | |
| Runs | 2,500,000 | 5,000,000 | 5,000,000 |
| Root age | Exponential(1) | Exponential(1) | |
| Origin time | | | Uniform(5,8) |
| log clock rate | Uniform (-5,10) | | |
| Clock rate | 10^log Clock Rate | | 10^log Clock Rate |
| Base rate | | Exponential(1) | |
| log Sigma | Normal(0,1) | Normal(0,1) | Normal(0,1) |
| Sigma | 10^log Sigma | 10^log Sigma | Exponential(1) |
| Tree prior | Uniform | Uniform | Fossilized Birth-Death Process |
| Speciation rate | | | Exponential(10) |
| Extinction rate | | | Beta(1.0,1.0) |
| Rho (probability of sampling extant lineage) | | | 1 |
| Psi (fossil sampling rate) | | | Exponential(10) |
| UCLN mean | | | Exponential(2.0) |
| UCLN Sigma | | | Exponential(3.0) |
| UCLN Mu | | | log(UCLN Mean)-(0.5*UCLN Variance) |
| UCLN Variance | | | UCLN Sigma^2 |
| Branch rates | | Base Rate*Gamma(2,2) | log Normal(UCLN Mu, UCLN Variance) |

[a]Any parameters not specifically mentioned in the text use suggested RevBayes priors

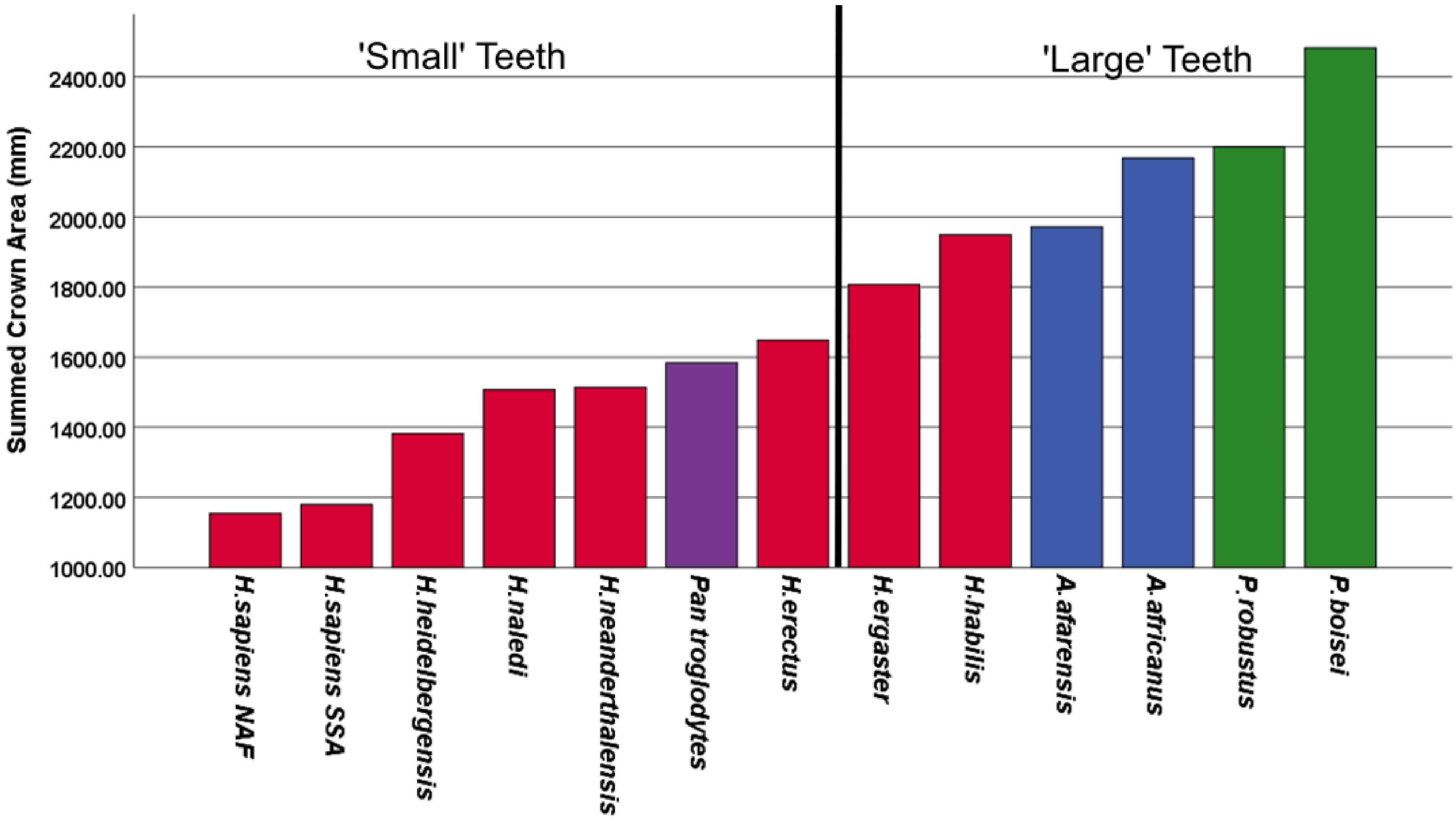


**SI Figure S1.** Bar graph of summed crown areas based on unscaled odontometric measurements [(MD x BL) x 16 total maxillary and mandibular teeth)] for all samples to illustrate dichotomy between 'large'- (1813-2483 total mm$^2$) and 'small'-toothed (1154-1648 mm$^2$) species responsible for influencing the UPGMA clusters in Figure 3 of the main text. Color-coding matches species in Figures 1, 2, and 5. See main text for details.

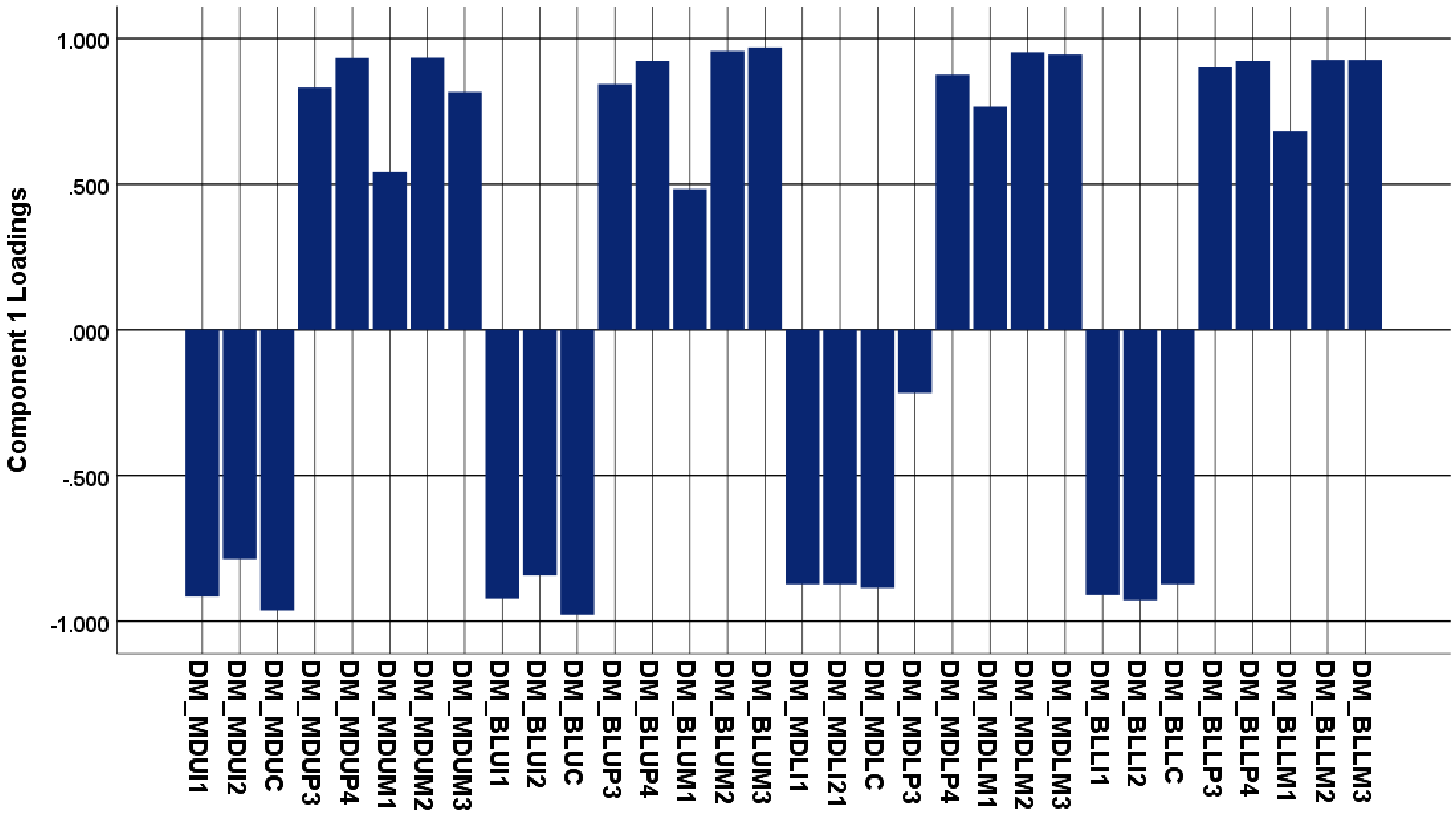


**SI Figure S2.** Graph of PCA loadings for the first component (74.6% of variance) from DM-corrected MD and BL dimensions of all teeth, from Table 3 in the main text. Bars illustrate those of most importance in driving sample location along the X-axis in Figure 5 of the main text. See the latter for details.

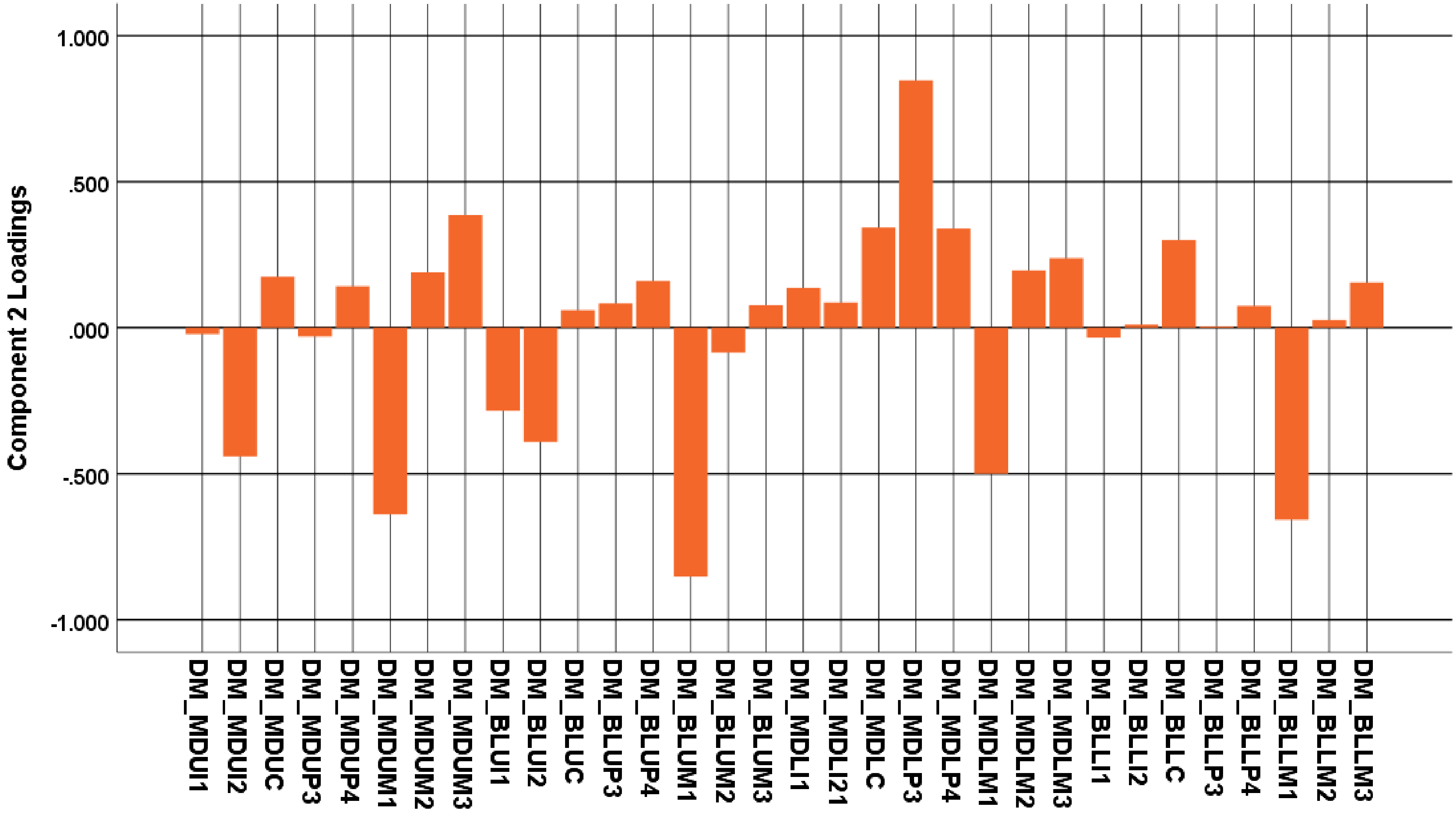


**SI Figure S3.** Graph of PCA loadings for the second component (11.6% of variance) from DM-corrected MD and BL dimensions of all teeth, from Table 3 in the main text. Bars illustrate those of most importance in driving sample location along the Y-axis in Figure 5 of the main text. See the latter for details.

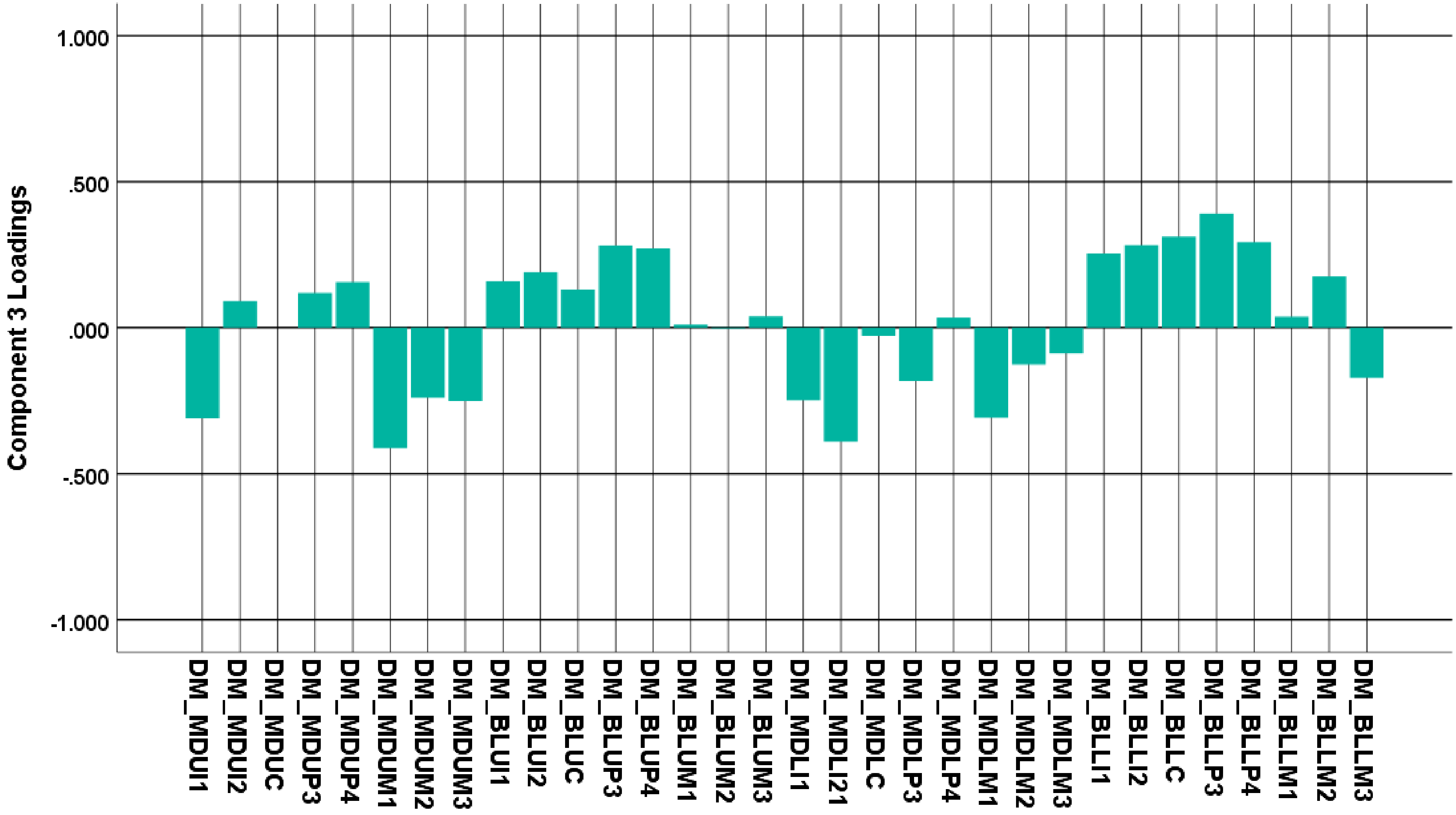


**SI Figure S4.** Graph of PCA loadings for the third component (4.6% of variance) from DM-corrected MD and BL dimensions of all teeth, from Table 3 in the main text. Bars illustrate those of most importance in driving sample location along the Z-axis in Figure 5 of the main text. See the latter for details.

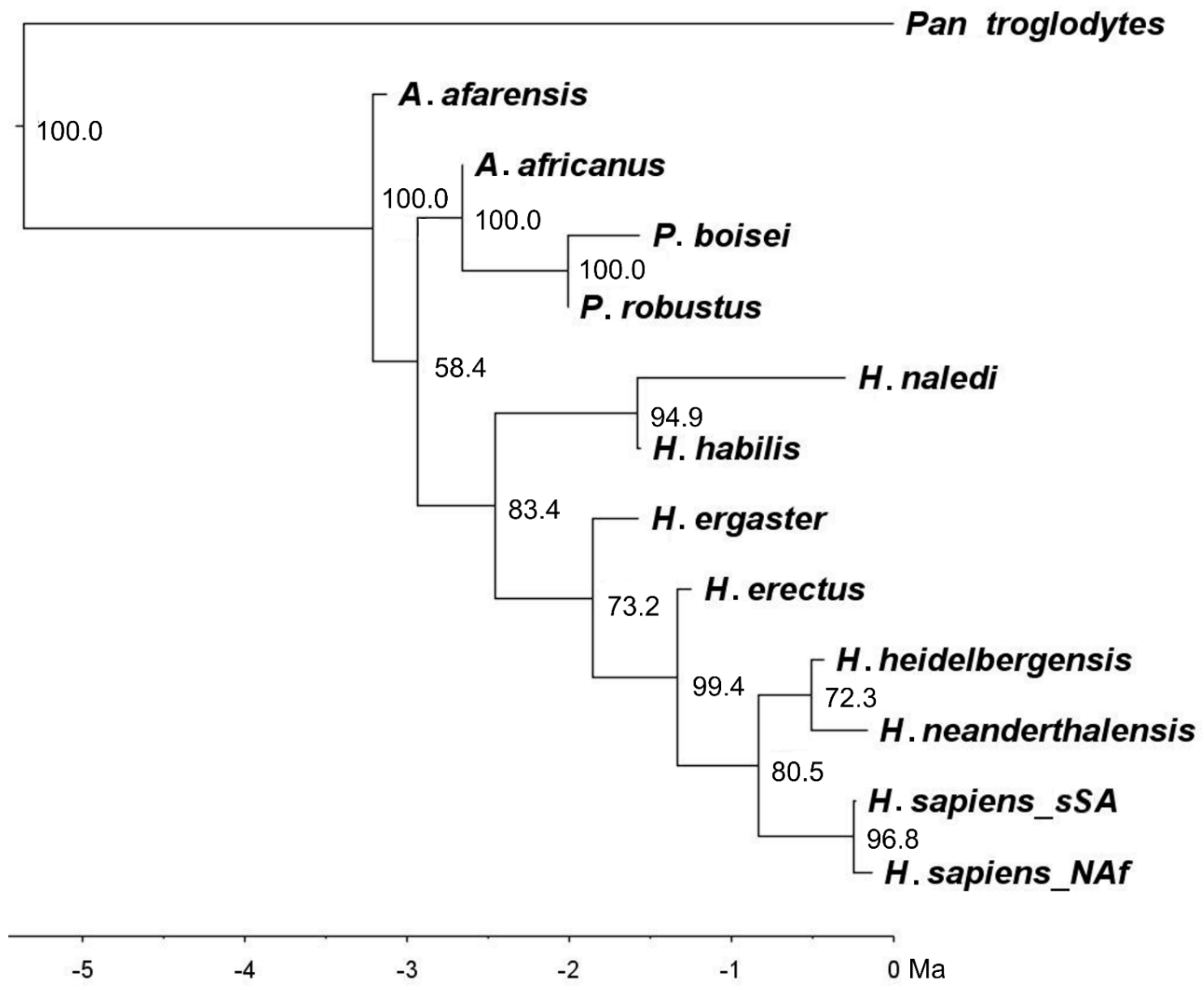


**SI Figure S5.** Bayesian inference phylogram (7,000,000 generations) from dated relaxed-clock FBD analysis with gap-weighted DM-scaled data under MKv model in MrBayes. Clade credibility values included. Rate variation is based on a gamma rates prior, but the clades match exactly with Figure 8. With a marginal likelihood of -595.12 it is the favored coded model. However, because is not directly comparable with the Bayesian FBD analysis of continuous data using a lognormal rates prior in RevBayes (see Materials and methods), it is not presented in the main text.

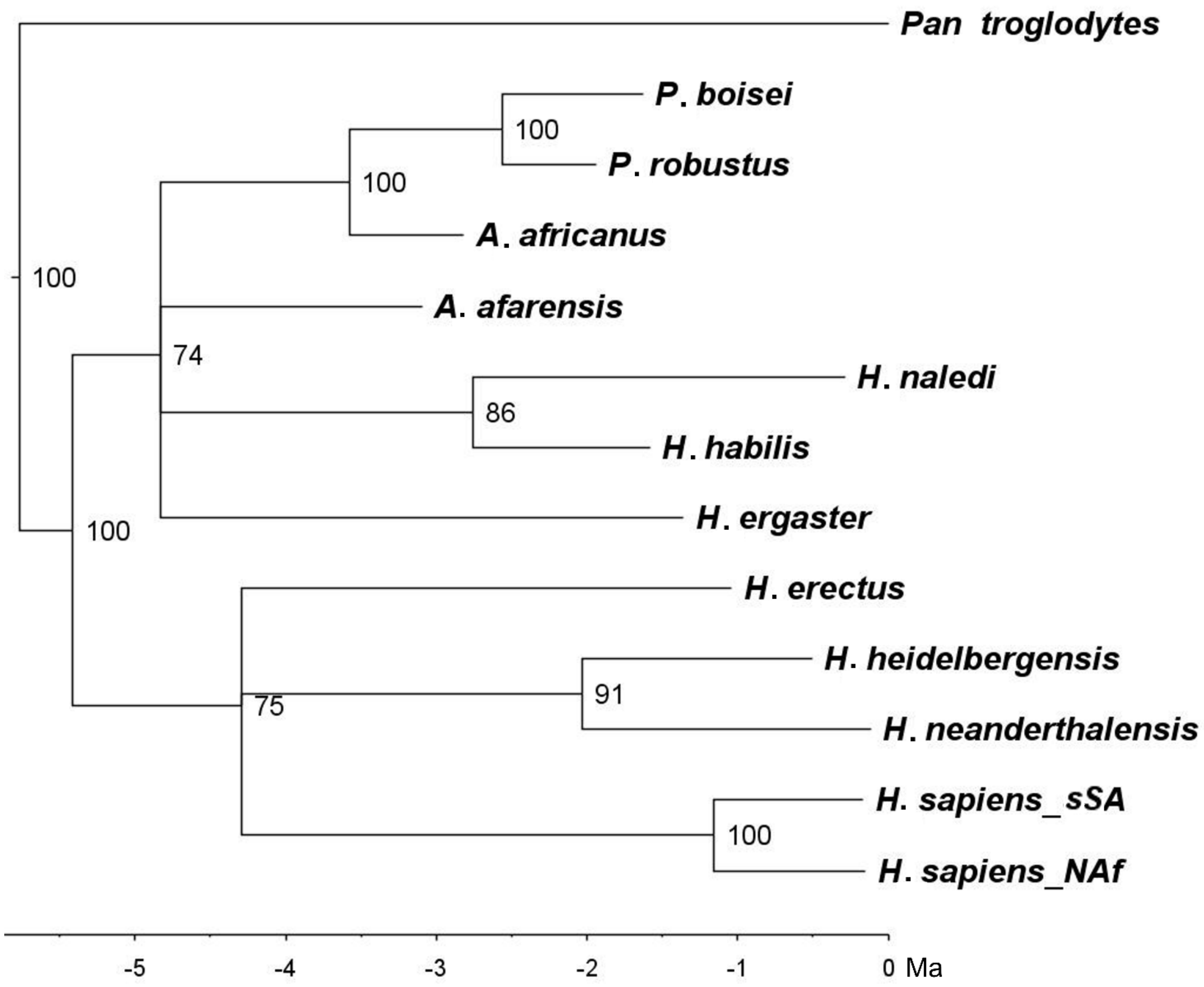


**SI Figure S6.** Bayesian inference phylogram (2,000,000 generations) from dated relaxed-clock analysis with gap-weighted DM-scaled data under MKv model in MrBayes. Clade credibility values included. Default equal rates (rather than gamma or lognormal) and uniform (rather than fossilized birth-death) branch lengths priors used for the model. Note two polytomies (c.f. Fig. 7) and non-representative branch lengths relative to divergence times based on fossil dates. See main text for details.

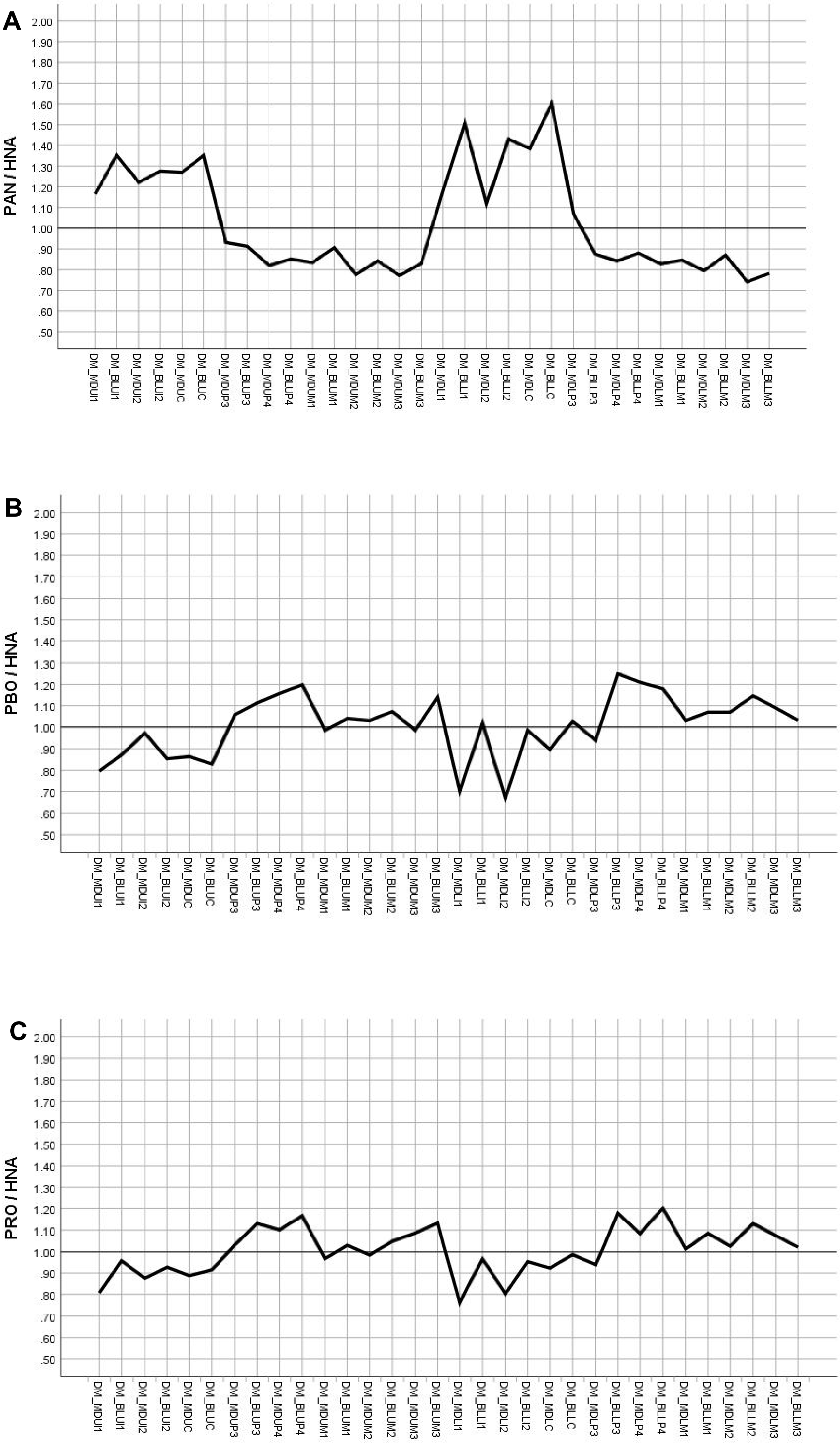
A
PAN / HNA
2.00
1.90
1.80
1.70
1.60
1.50
1.40
1.30
1.20
1.10
1.00
.90
.80
.70
.60
.50
DM_MDUI1
DM_BLUI1
DM_MDUI2
DM_BLUI2
DM_MDUC
DM_BLUC
DM_MDUP3
DM_BLUP3
DM_MDUP4
DM_BLUP4
DM_MDUM1
DM_BLUM1
DM_MDUM2
DM_BLUM2
DM_MDUM3
DM_BLUM3
DM_MDLI1
DM_BLLI1
DM_MDLI2
DM_BLLI2
DM_MDLC
DM_BLLC
DM_MDLP3
DM_BLLP3
DM_MDLP4
DM_BLLP4
DM_MDLM1
DM_BLLM1
DM_MDLM2
DM_BLLM2
DM_MDLM3
DM_BLLM3
B
PBO / HNA
2.00
1.90
1.80
1.70
1.60
1.50
1.40
1.30
1.20
1.10
1.00
.90
.80
.70
.60
.50
DM_MDUI1
DM_BLUI1
DM_MDUI2
DM_BLUI2
DM_MDUC
DM_BLUC
DM_MDUP3
DM_BLUP3
DM_MDUP4
DM_BLUP4
DM_MDUM1
DM_BLUM1
DM_MDUM2
DM_BLUM2
DM_MDUM3
DM_BLUM3
DM_MDLI1
DM_BLLI1
DM_MDLI2
DM_BLLI2
DM_MDLC
DM_BLLC
DM_MDLP3
DM_BLLP3
DM_MDLP4
DM_BLLP4
DM_MDLM1
DM_BLLM1
DM_MDLM2
DM_BLLM2
DM_MDLM3
DM_BLLM3
C
PRO / HNA
2.00
1.90
1.80
1.70
1.60
1.50
1.40
1.30
1.20
1.10
1.00
.90
.80
.70
.60
.50
DM_MDUI1
DM_BLUI1
DM_MDUI2
DM_BLUI2
DM_MDUC
DM_BLUC
DM_MDUP3
DM_BLUP3
DM_MDUP4
DM_BLUP4
DM_MDUM1
DM_BLUM1
DM_MDUM2
DM_BLUM2
DM_MDUM3
DM_BLUM3
DM_MDLI1
DM_BLLI1
DM_MDLI2
DM_BLLI2
DM_MDLC
DM_BLLC
DM_MDLP3
DM_BLLP3
DM_MDLP4
DM_BLLP4
DM_MDLM1
DM_BLLM1
DM_MDLM2
DM_BLLM2
DM_MDLM3
DM_BLLM3

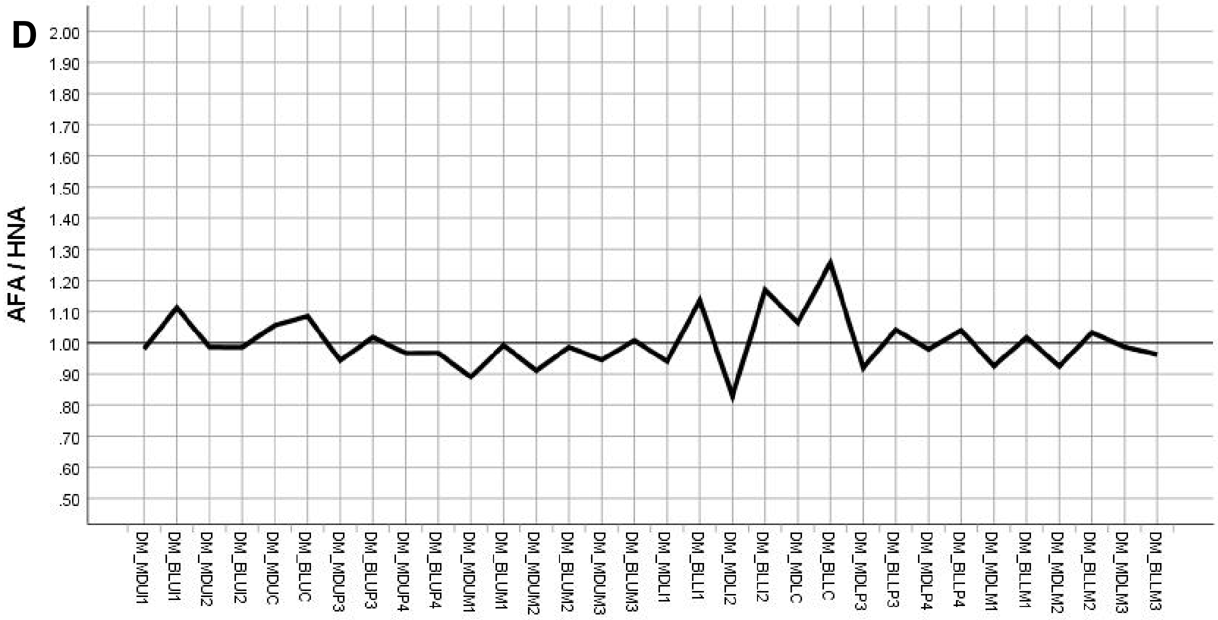
D
AFA / HNA
2.00
1.90
1.80
1.70
1.60
1.50
1.40
1.30
1.20
1.10
1.00
.90
.80
.70
.60
.50
DM_MDUI1
DM_BLUI1
DM_MDUI2
DM_BLUI2
DM_MDUC
DM_BLUC
DM_MDUP3
DM_BLUP3
DM_MDUP4
DM_BLUP4
DM_MDUM1
DM_BLUM1
DM_MDUM2
DM_BLUM2
DM_MDUM3
DM_BLUM3
DM_MDLI1
DM_BLLI1
DM_MDLI2
DM_BLLI2
DM_MDLC
DM_BLLC
DM_MDLP3
DM_BLLP3
DM_MDLP4
DM_BLLP4
DM_MDLM1
DM_BLLM1
DM_MDLM2
DM_BLLM2
DM_MDLM3
DM_BLLM3

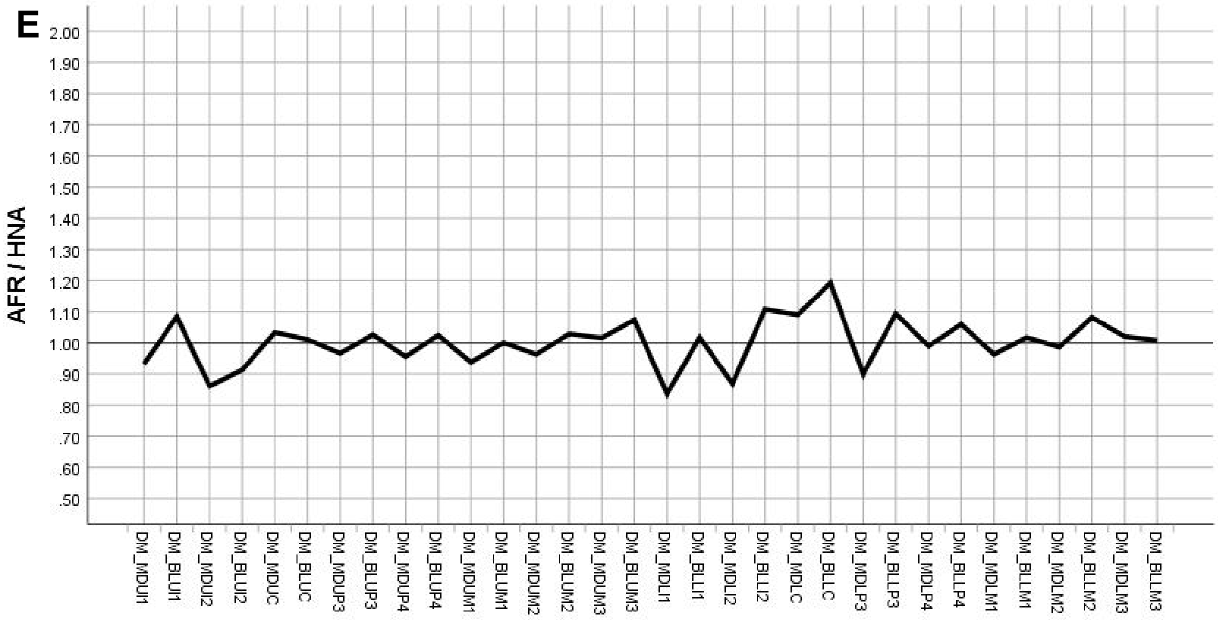
E
AFR / HNA
2.00
1.90
1.80
1.70
1.60
1.50
1.40
1.30
1.20
1.10
1.00
.90
.80
.70
.60
.50
DM_MDUI1
DM_BLUI1
DM_MDUI2
DM_BLUI2
DM_MDUC
DM_BLUC
DM_MDUP3
DM_BLUP3
DM_MDUP4
DM_BLUP4
DM_MDUM1
DM_BLUM1
DM_MDUM2
DM_BLUM2
DM_MDUM3
DM_BLUM3
DM_MDLI1
DM_BLLI1
DM_MDLI2
DM_BLLI2
DM_MDLC
DM_BLLC
DM_MDLP3
DM_BLLP3
DM_MDLP4
DM_BLLP4
DM_MDLM1
DM_BLLM1
DM_MDLM2
DM_BLLM2
DM_MDLM3
DM_BLLM3

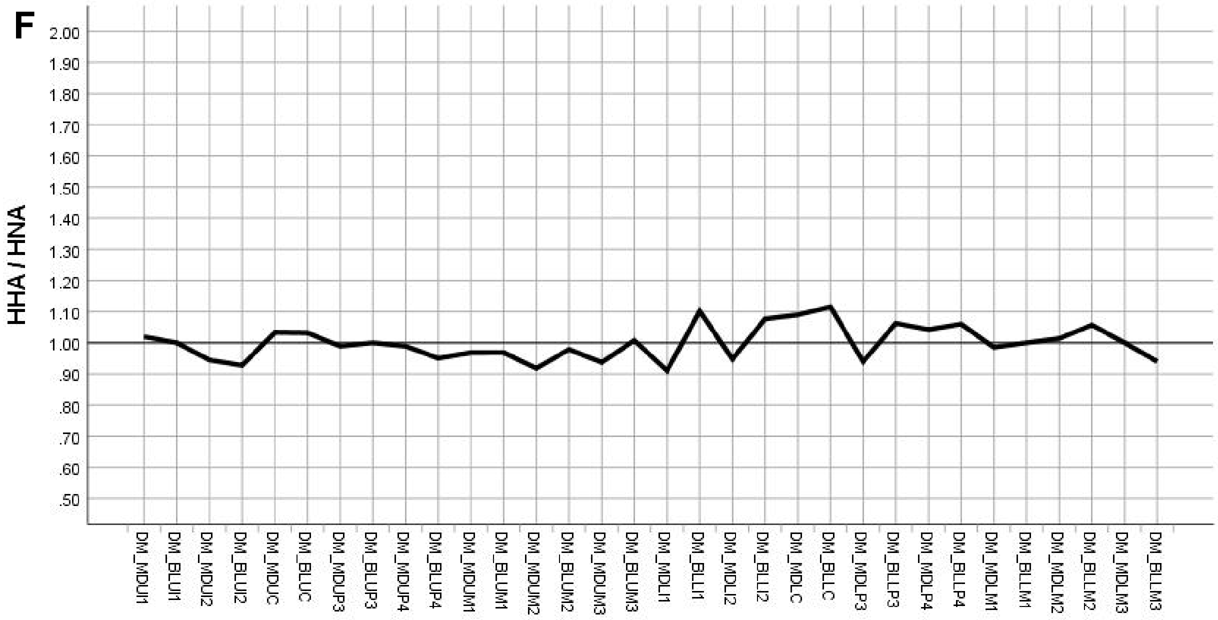
F
HHA / HNA
2.00
1.90
1.80
1.70
1.60
1.50
1.40
1.30
1.20
1.10
1.00
.90
.80
.70
.60
.50
DM_MDUI1
DM_BLUI1
DM_MDUI2
DM_BLUI2
DM_MDUC
DM_BLUC
DM_MDUP3
DM_BLUP3
DM_MDUP4
DM_BLUP4
DM_MDUM1
DM_BLUM1
DM_MDUM2
DM_BLUM2
DM_MDUM3
DM_BLUM3
DM_MDLI1
DM_BLLI1
DM_MDLI2
DM_BLLI2
DM_MDLC
DM_BLLC
DM_MDLP3
DM_BLLP3
DM_MDLP4
DM_BLLP4
DM_MDLM1
DM_BLLM1
DM_MDLM2
DM_BLLM2
DM_MDLM3
DM_BLLM3

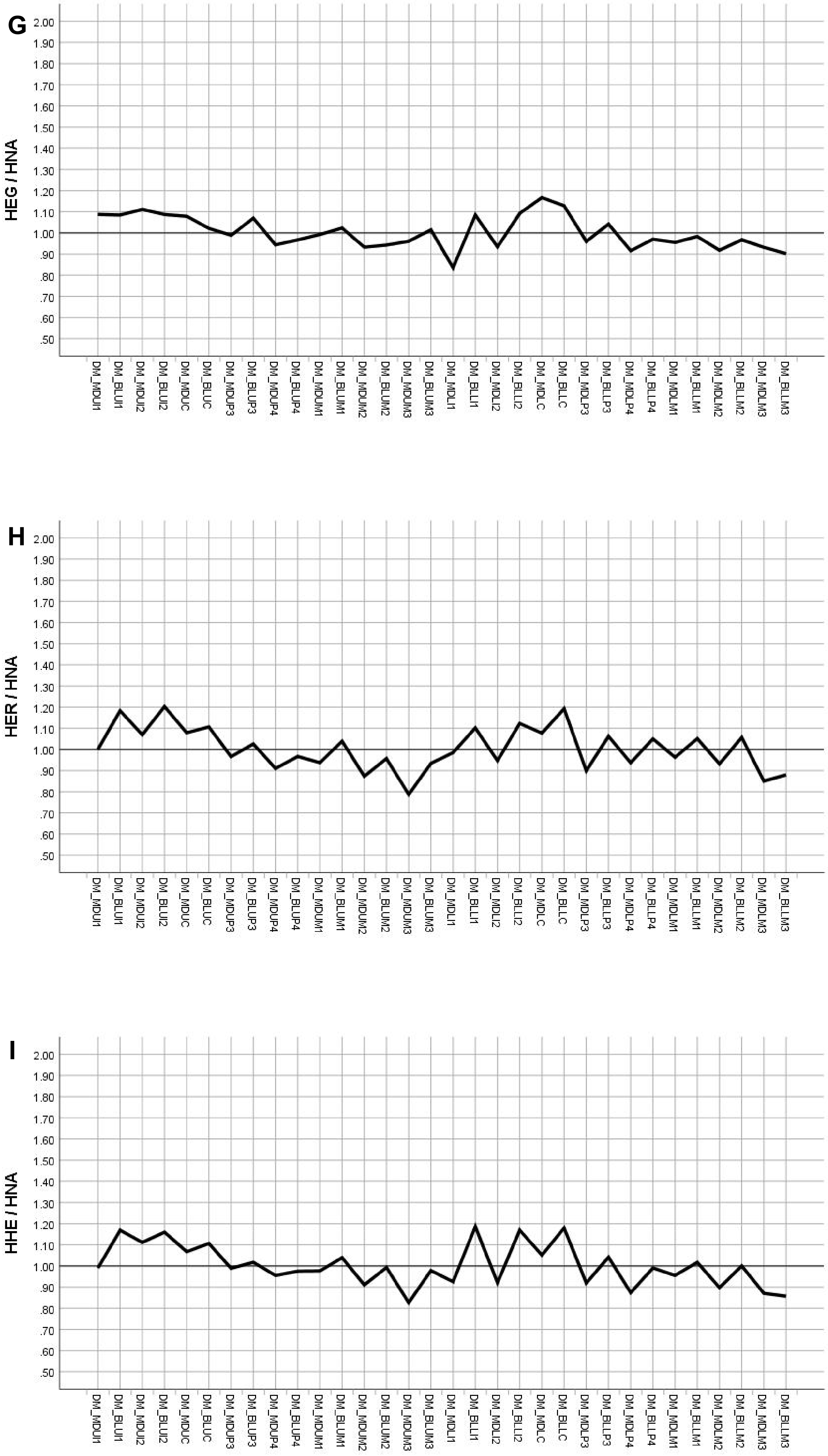
G
HEG / HNA
H
HER / HNA
I
HHE / HNA
2.00
1.90
1.80
1.70
1.60
1.50
1.40
1.30
1.20
1.10
1.00
.90
.80
.70
.60
.50
DM_MDUI1
DM_BLUI1
DM_MDUI2
DM_BLUI2
DM_MDUC
DM_BLUC
DM_MDUP3
DM_BLUP3
DM_MDUP4
DM_BLUP4
DM_MDUM1
DM_BLUM1
DM_MDUM2
DM_BLUM2
DM_MDUM3
DM_BLUM3
DM_MDLI1
DM_BLLI1
DM_MDLI2
DM_BLLI2
DM_MDLC
DM_BLLC
DM_MDLP3
DM_BLLP3
DM_MDLP4
DM_BLLP4
DM_MDLM1
DM_BLLM1
DM_MDLM2
DM_BLLM2
DM_MDLM3
DM_BLLM3

J

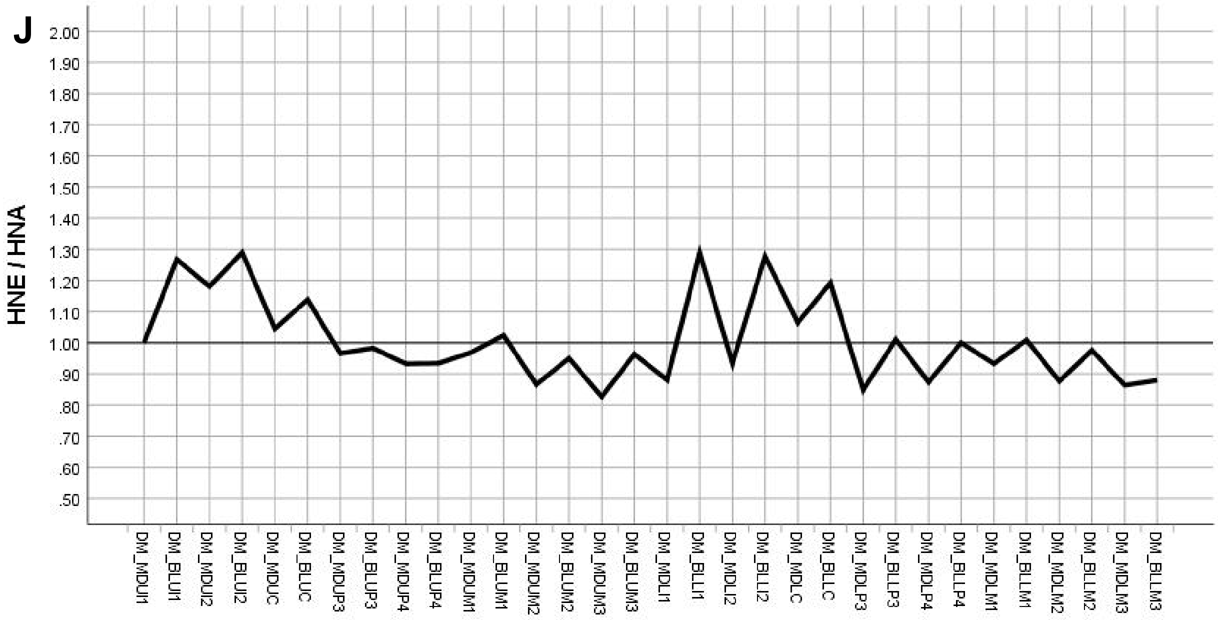


K

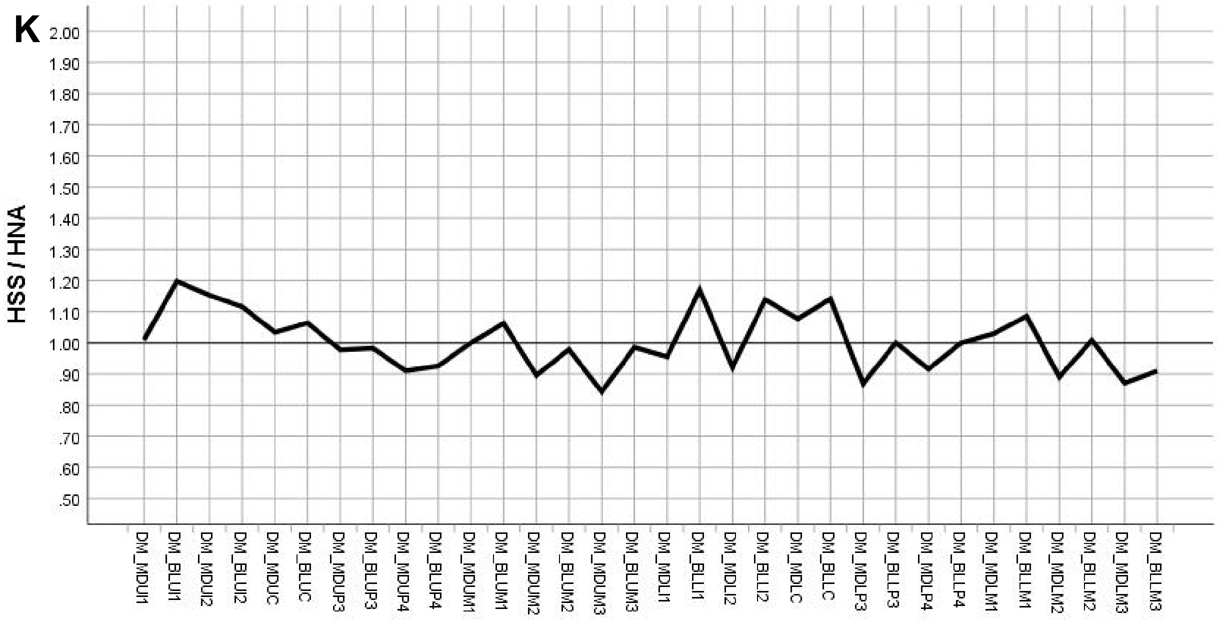


L

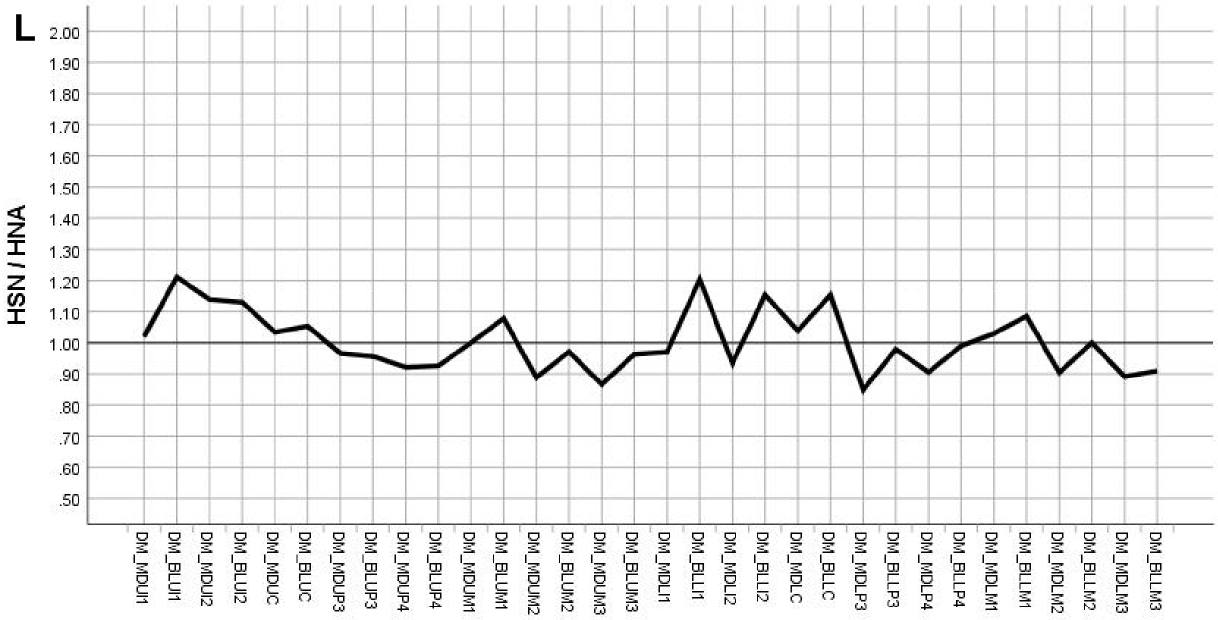


**SI Figure S7.** Scatterplots of quotients between the dividends A) PAN, B) PBO, C) PRO, D) AFA, E) AFR, F) HHA, G) HEG, H) HER, I) HHE, J) HNE, K) HSS, L) HSN and the divisor HNA for the DM-scaled data, to visualize key characters in *H. naledi* relative to each comparative sample. Note closest similarity in F, between *H. habilis* and *H. naledi.*
